# LIN28B controls the regenerative capacity of neonatal murine auditory supporting cells through activation of mTOR signaling

**DOI:** 10.1101/2020.05.31.126193

**Authors:** Xiaojun Li, Angelika Doetzlhofer

**Affiliations:** The Solomon H. Snyder Department of Neuroscience, Center for Sensory Biology, The Johns Hopkins University School of Medicine, Baltimore, Maryland 21205, USA

**Keywords:** hair cell regeneration, supporting cell, de-differentiation, inner ear cochlea, LIN28A, LIN28B, Let-7 miRNA, mTOR pathway

## Abstract

Mechano-sensory hair cells within the inner ear cochlea are essential for the detection of sound. In mammals, cochlear hair cells are only produced during development and their loss, due to disease or trauma, is a leading cause of deafness. In the immature cochlea, prior to the onset of hearing, hair cell loss stimulates neighboring supporting cells to act as hair cell progenitors and produce new hair cells. However, for reasons unknown, such regenerative capacity (plasticity) is lost once supporting cells undergo maturation. Here, we demonstrate that the RNA binding protein LIN28B plays an important role in the production of hair cells by supporting cells and provide evidence that the developmental drop in supporting cell plasticity in the mammalian cochlea is, at least in part, a product of declining LIN28B-mTOR activity. Employing murine cochlear organoid and explant cultures to model mitotic and non-mitotic mechanisms of hair cell generation, we show that loss of *Lin28b* function, due to its conditional deletion, or due to overexpression of the antagonistic miRNA *let-7g*, suppressed Akt-mTORC1 activity and renders young, immature supporting cells incapable of generating hair cells. Conversely, we found that LIN28B overexpression increased Akt-mTORC1 activity and allowed supporting cells that were undergoing maturation to de-differentiate into progenitor-like cells and to produce hair cells via mitotic and non-mitotic mechanisms. Finally, using the mTORC1 inhibitor rapamycin, we demonstrate that LIN28B promotes supporting cell plasticity in an mTORC1-dependent manner.

**SIGNIFICANCE STATEMENT:** Cochlear hair cell loss is a leading cause of deafness in humans and other mammals. In the immature cochlea lost hair cells are regenerated by neighboring glia-like supporting cells. However, for reasons unknown, such regenerative capacity is rapidly lost as supporting cells undergo maturation. Here we identify a direct link between LIN28B-mTOR activity and supporting cell plasticity. Mimicking later developmental stages, we found that loss of the RNA binding protein LIN28B attenuated mTOR signaling and rendered young, immature supporting cells incapable of producing hair cells. Conversely, we found that re-expression of LIN28B reinstated the ability of maturing supporting cells to revert to a progenitor-like state and generate hair cells via activation of mTOR signaling.

---

The cochlea, located in the inner ear, contains highly specialized mechano-receptor cells, termed hair cells, which are critical for our ability to detect sound. In mammals, auditory hair cells are only produced during embryonic development and hair cell loss due to aging, disease or trauma is a leading cause for hearing impairment and deafness in humans. Non-mammalian vertebrates, such as birds, are capable of regenerating hair cells within their auditory and vestibular sensory organs throughout their lifetime (reviewed in (1)). The source of the newly generated (regenerated) hair cells are glia-like supporting cells. Sharing a close lineage relationship, hair cells and supporting cells originate from a common pool of progenitor cells, termed pro-sensory cells (2, 3). The transcription factor SOX2 is essential for the establishment and maintenance of pro-sensory cells (4) and plays a critical role in the induction of *Atoh1* during hair cell formation (5, 6). ATOH1, a bHLH transcriptional activator, is required for hair cell formation in the developing inner ear (7) and its reactivation in supporting cells is an essential step in the process of hair cell regeneration (8, 9).

In birds, loss of auditory hair cells induces adjacent supporting cells to either directly convert (trans-differentiate) into hair cells (10, 11), or alternatively, to re-enter the cell cycle and after rounds of cell division to produce both hair cells and supporting cells (12, 13). In response to hair cell death, cochlear supporting cells in newborn mice have been recently shown to re-enter the cell cycle and produce new hair cells (14, 15). The injury-induced regenerative response in the neonatal/ early postnatal cochlea can be greatly enhanced by genetic or pharmacologic inhibition of Notch signaling (16-18) and/or over-activation of the Wnt/β-catenin signaling (19-22). However, mice are born deaf and their cochlear hair cells and supporting cells are not functional (mature) until the onset of hearing at P12-P13 (23-25). Recent studies uncovered that as early as postnatal day 5/6 (P5/P6) murine cochlear supporting cells fail to regenerate hair cells in response to injury (14), inhibition of Notch signaling (26), or over-activation of Wnt/β-catenin signaling (27). What causes the rapid decline in supporting cell plasticity during the first postnatal week is currently unknown. We previously demonstrated that a regulatory circuit consisting of LIN28B and *let-7* miRNAs modulates the production of new hair cells in stage P2 murine cochlear explants, with LIN28B promoting new hair cell production and *let-7* miRNAs suppressing it (28). The closely related RNA binding proteins LIN28A and LIN28B (LIN28A/B) and members of the *let-7* family of miRNAs belong to an evolutionary highly conserved network of genes, initially identified in *C-elegans* for their role in developmental timing (heterochrony)(29, 30). *Let-7* miRNAs and LIN28A/B are mutual antagonists, which repress each other’s expression. LIN28A/B inhibit the biogenesis of *let-7* miRNAs through direct binding to primary and precursor-*let-7* transcripts. In turn, *let-7* miRNAs interfere with the translation of *Lin28a* and *Lin28b* mRNAs by binding to their 3’UTR (reviewed in (31)). While *let-7* miRNAs are linked to a post-mitotic, terminal differentiated state, LIN28A/B are positive regulators of stemness, organismal growth and metabolism (reviewed in (32)). How LIN28B/*let-7* influence the production of hair cells by supporting cells and whether the LIN28B/*let-7* circuit plays a role in the developmental decline of supporting cell plasticity is currently unknown.

Here, we provide evidence that the decline in supporting cell plasticity in the murine cochlea is, at least in part, due to diminished LIN28B-mTORC1 activity. Using organoid and explant cultures to model mitotic and non-mitotic hair cell production by supporting cells, we show that diminished LIN28B-mTOR activity, due to *let-7g* overexpression or targeted deletion of *Lin28a/b*, accelerated the developmental decline in supporting cell plasticity and rendered, otherwise plastic, immature supporting cells incapable of producing new hair cells. Finally, we provide evidence that LIN28B stimulates mitotic and non-mitotic hair cell production via Akt-mTORC1 activation. We found that rapamycin, a selective inhibitor of the mTORC1 kinase complex, blocked LIN28B-induced hair cell production in cochlear explant and organoid culture.

## RESULTS

### The ability of murine cochlear epithelial cells to form and grow hair cell containing organoids sharply declines within the first postnatal week

Cochlear supporting cells, isolated from newborn mice, re-enter the cell cycle and produce hair cells when cultured on a feeder layer of periotic mesenchymal cells in the presence of mitogens. However, mature cochlear supporting cells isolated from mice at stage P14 fail to proliferate and produced only few scattered hair cells when cultured under identical conditions (33). A similar decline in the potential to proliferate and self-renew is observed when cochlear epithelial cells, including supporting cells, are propagated using neurosphere culture conditions (34). To address the underlying molecular mechanisms that cause the decline in supporting cell plasticity, we employed a recently developed 3D organoid system, which allows Wnt-responsive cochlear epithelial cells (supporting cells) to propagate and differentiate into large hair cell –containing organoids (35).

Employing this novel organoid culture system (Fig. 1 *A*, expansion), we found that cochlear epithelial cells from mice, stage P2 (P2 organoid culture) readily formed large organoids (Fig. 1 *B*, P2). By contrast, cochlear epithelial cells from mice, stage P5 (P5 organoid culture) showed a greatly diminished capacity to form and grow organoids (Fig. 1 *B*, P5). After 5 days of expansion, P5 organoid cultures contained 3-times fewer organoids than P2 organoid cultures (Fig. 1 *B* and *C*) and on average, organoids in P5 organoid culture were only a quarter the size of organoids in P2 organoid cultures (Fig. 1 *B* and *D*). To address potential defects in cell proliferation, P2 and P5 organoid cultures received a 1-hour pulse of EdU after which organoids were harvested and co-stained for SOX2. Prior to cochlear differentiation, SOX2 expression identifies pro-sensory cells and later marks supporting cells and Kölliker’s cells, a transient population of supporting-like cells located at the medial border of the sensory epithelium. In addition, SOX2 is transiently expressed in nascent hair cells (6, 36). We found that SOX2^+^ cells in P5 organoids proliferated at a 25% lower rate (% of EdU+ cells) than SOX2^+^ cells in P2 organoids (Fig. 1 *E* and *F*), indicating that the observed defects in P5 organoid formation and growth were the product of reduced cell proliferation.

**Fig.1.**
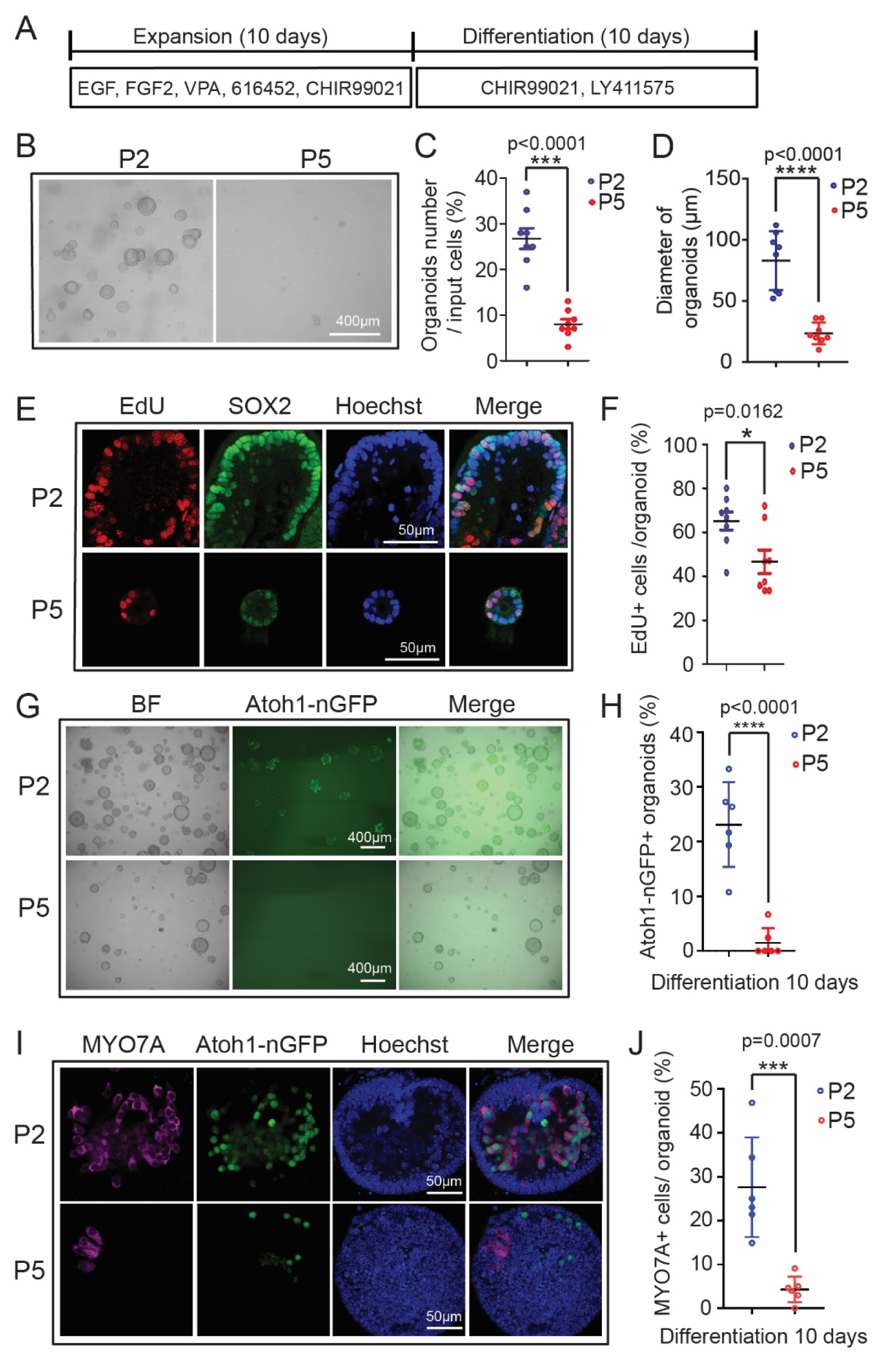
Cochlear epithelial cells from stage P5 mouse pups fail to expand and produce hair cells in organoid culture. Organoid cultures were established from cochlear epithelia cells obtained from *Atoh1-nGFP* transgenic mice, stages postnatal day 2 (P2) (P2 organoid culture) and P5 (P5 organoid culture). Atoh1-nGFP reporter expression marks nascent hair cells. Graphed are individual data points and mean ± SD. 2-tailed, unpaired Student’s t-test was used to calculate p-values. All shown data is from 2 independent experiments. (*A*) Experimental design. (*B*) Bright field (BF) images of P2 and P5 cochlear organoid cultures after 5 days of expansion. (*C*) Organoid forming efficiency in P2 (blue) and P5 (red) cultures (*A*) (n=8 animals per group). (*D*) Organoid diameters in P2 (blue) and P5 (red) cultures in (*A*) (n=8 animals per group). (*E*) Cell proliferation in P2 and P5 organoids. A single EdU pulse was given at 5 days of expansion and EdU incorporation (red) was analyzed 1 hour later. SOX2 (green) marks supporting cells/ pro-sensory cells and Hoechst (blue) labels cell nuclei. (*F*) Percentage of EdU+ cells in P2 and P5 organoids in (*E*) (n=8 animals per group). (*G*) BF and green fluorescent images (Atoh1-nGFP) of P2 and P5 organoid cultures after 10 days of differentiation. (*H*) Percentage of Atoh1-nGFP^+^ in P2 and P5 organoids in (*G*) (n=6 animals per group). (*I*) Confocal images of P2 and P5 organoid cultures after 10 days of differentiation. Newly formed hair cells express Atoh1-nGFP (green) and MYO7A (magenta). (*J*) Percentage of MYO7A^+^ hair cells per organoid in (*I*) (n=6 animals per group). Note that the individual data points in *D, F* and *J* represent the average values per animal.

Next, we analyzed the ability of P2 and P5 organoids to produce hair cells (Fig.1 *A*, differentiation). Hair cell formation in P2 and P5 organoid cultures was monitored using Atoh1-nGFP reporter expression (37). *Atoh1* (Atoh1-nGFP) expression is high in nascent hair cells, but absent in hair cells that underwent maturation, which allows to distinguish between existing and nascent hair cells (37, 38). After 10 days of differentiation, approximately 25% of organoids in P2 organoid cultures contained Atoh1-nGFP^+^ cells (Fig. 1 *G* and *H*, P2), which formed large clusters and co-expressed the hair cell-specific protein myosin VIIa (MYO7A) (Fig. 1 *I* and *J*, P2). By contrast, less than 2% of organoids in P5 organoid cultures contained few, scattered Atoh1-nGFP^+^ cells and largely lacked myosinVIIa expression (Fig. 1 *G*-*J*, P5). The observed decline in supporting cell plasticity between P2 and P5 correlated with a drop in *Lin28b* mRNA and LIN28B protein expression within cochlear epithelial cells *in vivo* (Fig. S1 *A* and *B*).

### LIN28B re-activation re-instates cochlear supporting cell plasticity

In order to address whether there is a link between LIN28B protein levels and supporting cell plasticity, we re-expressed LIN28B in stage P5 cochlear organoids using *iLIN28B* transgenic mice. In this double transgenic mouse model, a flag-tagged human *LIN28B* transgene is expressed under the control of a TRE promoter (*Col1a-TRE-LIN28B*)(39), which in the presence of ubiquitously expressed *R26-rtTA-M2* transgene and doxycycline (dox) allows for robust induction of LIN28B protein (28, 39). First, we expanded cochlear epithelia cells from P5 *iLIN28B* transgenic mice and control littermates (lack *LIN28B* transgene) in the presence of dox for 13 days (Fig. 2 *A*). We found that LIN28B overexpression increased the efficiency of organoid formation by more than 5-fold, as well as doubled the average organoid size (diameter) compared to control (Fig. 2 *B*-*D*). The increase in organoid size in response to LIN28B overexpression was accompanied by a 2-fold higher rate of cell proliferation during the early phase of expansion (Fig. 2 *F* and *G*). Coinciding with the peak of cell proliferation, *Sox2* transcript levels in LIN28B overexpressing organoids transiently decreased more than 3-fold compared to control organoids (Fig. 2 *E*) and the majority of proliferating cells (EdU^+^) in LIN28B overexpressing organoids expressed SOX2 at a low level (Fig. 2 *H*, SOX2^low^, EdU^+^).

**Fig.2.**
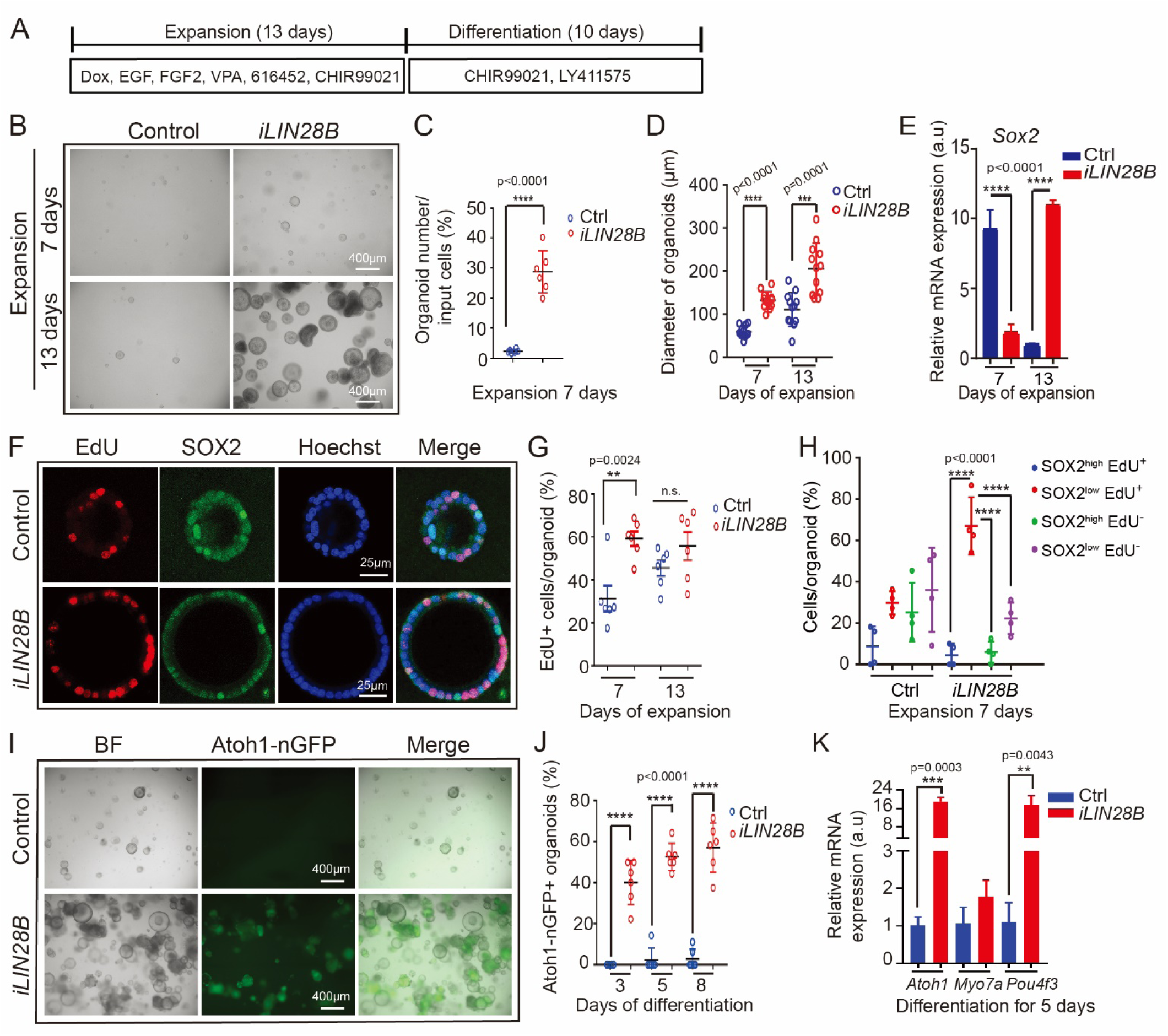
LIN28B overexpression promotes cochlear epithelial cell expansion and hair cell production. Cochlear organoid cultures were established from stage P5 *Atoh1-nGFP; iLIN28B* transgenic mice and control littermates that lacked the *LIN28B* transgene. Culture media were replenished every other day. Atoh1-GFP reporter labeled nascent hair cells. Bars in *(E)* and *(K)* represent mean± SD, otherwise individual data points and their mean ± SD were plotted. 2-tailed, unpaired Student’s t-test was used to calculate p-values. (*A*) Experimental strategy. *(B)* BF images of control and LIN28B overexpressing (*iLIN28B*) organoid cultures at 7 and 13 days of expansion. (*C*) Organoid forming efficiency in control (Ctrl, blue) and LIN28B (*iLIN28B*, red) overexpressing cultures (n=6 animals per group, from 2 independent experiments). (*D*) Organoid diameters in control (Ctrl, blue) and LIN28B (*iLIN28B*, red) overexpressing cultures after 7 and 13 days of expansion (n=12 animals per group). *(E)* RT-qPCR-based analysis of *Sox2* mRNA expression in control (Ctrl, blue bar) and LIN28B overexpressing (iLIN28B, red bar) organoids at 7 and 13 days of expansion (n=3 animals per group, from 1 representative experiment, 3 independent experiments). (*F*) Cell proliferation in control and LIN28B overexpressing organoids. A single EdU pulse was given at 7 days (shown) or 13 days of expansion and EdU incorporation (red) was analyzed 2 hours later. SOX2 (green) marks supporting cells/ pro-sensory cells, Hoechst labels cell nuclei (blue). (*G*) Percentage of EdU^+^ cells in control (Ctrl, blue) and LIN28B overexpressing (*iLIN28B*, red) organoids at 7 and 13 days of expansion (n=6 animals per group, from 2 independent experiments, n.s. not significant). (*H*) EdU incorporation in SOX2-high and SOX2-low expressing cells (n=4 animals per group, from 2 independent experiments). Note that the majority of EdU+ cells in LIN28B overexpressing organoids expressed SOX2 at a low level (red, SOX2^low^ EdU^+^). *(I)* BF and green fluorescent (Atoh1-nGFP) images of control and LIN28B overexpressing organoids after 8 days of differentiation. (*J*) Percentage of Atoh1-nGFP^+^ organoids in control (Ctrl) and LIN28B overexpressing (*iLIN28B*) cultures after 3, 5 and 8 days of differentiation (n=6 animals per group, from 2 independent experiments). *(K)* RT-PCR of hair cell-specific genes (*Atoh1, Myo7a, Pou4f3*) in control (Ctrl, blue bar) and LIN28B overexpressing (*iLIN28B*, red bar) organoids after 5 days of differentiation (n=3 animals per group, from 1 representative experiment, 3 independent experiments). Note that the individual data points in (*D*), (*G*) and (*H*) represent the average values per animal. Abbreviations: dox, doxycycline; a.u., arbitrary unit.

Next, we analyzed the capacity of control and LIN28B overexpressing organoids to produce hair cells. We found that after 3 days of differentiation, about 40% of LIN28B overexpressing organoids contained Atoh1-nGFP^+^ cell clusters, which increased to nearly 60% after 8 days of differentiation (Fig. 2 *I* and *J, iLIN28B*). By contrast, less than 5% of control organoids contained Atoh1-nGFP^+^ cells after 8 days of differentiation (Fig. 2 *I* and *J*, control). Confirming the presence of hair cells in LIN28B overexpressing organoids, we found that hair cell-specific transcripts (*Atoh1, Pou4f3)* were more than 10-fold up-regulated in LIN28B overexpressing organoids compared to control organoids (Fig. 2 *K*). Mimicking hair cell development *in vivo*, nascent Atoh1-nGFP^+^ hair cells first upregulated myosin VIIa (MYO7A)(Fig. S *B*), followed by the upregulation of calretinin, a protein enriched in the inner hair cells (40)(Fig. S *A, C, D*). After 7 days of differentiation, Atoh1-nGFP^+^ hair cells also started to up-regulate prestin, a protein selectively expressed in outer hair cells (Fig. S2 *F*) and LIN28B overexpressing organoids upregulated the expression of oncomodulin (*Ocm*), an outer hair cell-specific gene (41) and *Fgf8*, an inner hair cell-specific gene (42) (Fig. S2 *G*). *In vivo*, cochlear hair cells only transiently express *Atoh1* (Atoh1-nGFP) and SOX2 and their expression is lost once cochlear hair cells undergo maturation (6, 43). Similarly, we found that while MYO7A+ hair cells initially co-expressed Atoh1-nGFP and SOX2 in LIN28B overexpressing organoids (Fig. S2 *E*, Diff. 5 days), at later stages of culture MYO7A+ hair cells that lacked Atoh1-nGFP and SOX2 expression were observed (Fig. S2 *E*, Diff. 10 days). Together, these findings indicate that the newly formed hair cells in LIN28B overexpressing organoids acquired mature characteristics and specialized into inner and outer hair cell-like cells.

To be able to monitor the behavior of supporting cells within control and LIN28B overexpressing organoids, we generated control and *iLIN28B* transgenic mice that carried the *p27-GFP* BAC transgene. GFP expression in *p27-GFP* transgenic mice is under the control of the p27/Kip1 (*cdkn1b*) gene locus (44). P27/Kip1, a cyclin dependent kinase inhibitor, is essential for forcing pro-sensory cells out of the cell cycle and for maintaining supporting cells in a post-mitotic state (45-47). In the cochlear sensory epithelium p27-GFP is selectively expressed in supporting cells. Other cell types including hair cells don’t express p27-GFP or, in case of Kölliker’s cells, express p27-GFP at a much lower level than supporting cells (33). Importantly, p27-GFP expression is rapidly lost when supporting cells re-enter the cell cycle (33), allowing us to pinpoint the peak of cell-cycle re-entry of supporting cells in cochlear organoids. There was no significant difference in the number of p27-GFP+ organoids in control and LIN28B overexpressing cultures during early expansion (5 days), suggesting that supporting cells survived equally well in control and LIN28B overexpressing cultures. In control cultures the percentage of p27-GFP^+^ organoids declined during the remainder of the culture and after 7 days of differentiation less than 10% of organoids contained p27-GFP^+^ supporting cells (Fig. 3 *A* and *B*, Ctrl, blue line), suggesting that the vast majority of supporting cells in control organoids failed to re-enter the cell cycle. By contrast, p27-GFP expression in LIN28B overexpressing organoids, which during the latter half of expansion was near absent, dramatically increased upon differentiation. After 7 days of differentiation close to 50% of organoids in LIN28B overexpressing cultures contained large clusters of p27-GFP^+^ cells, indicating that LIN28B overexpression enabled stage P5 supporting cells to propagate in organoid culture (Fig. 3 *A* and *B, iLIN28B* red line).

**Fig.3.**
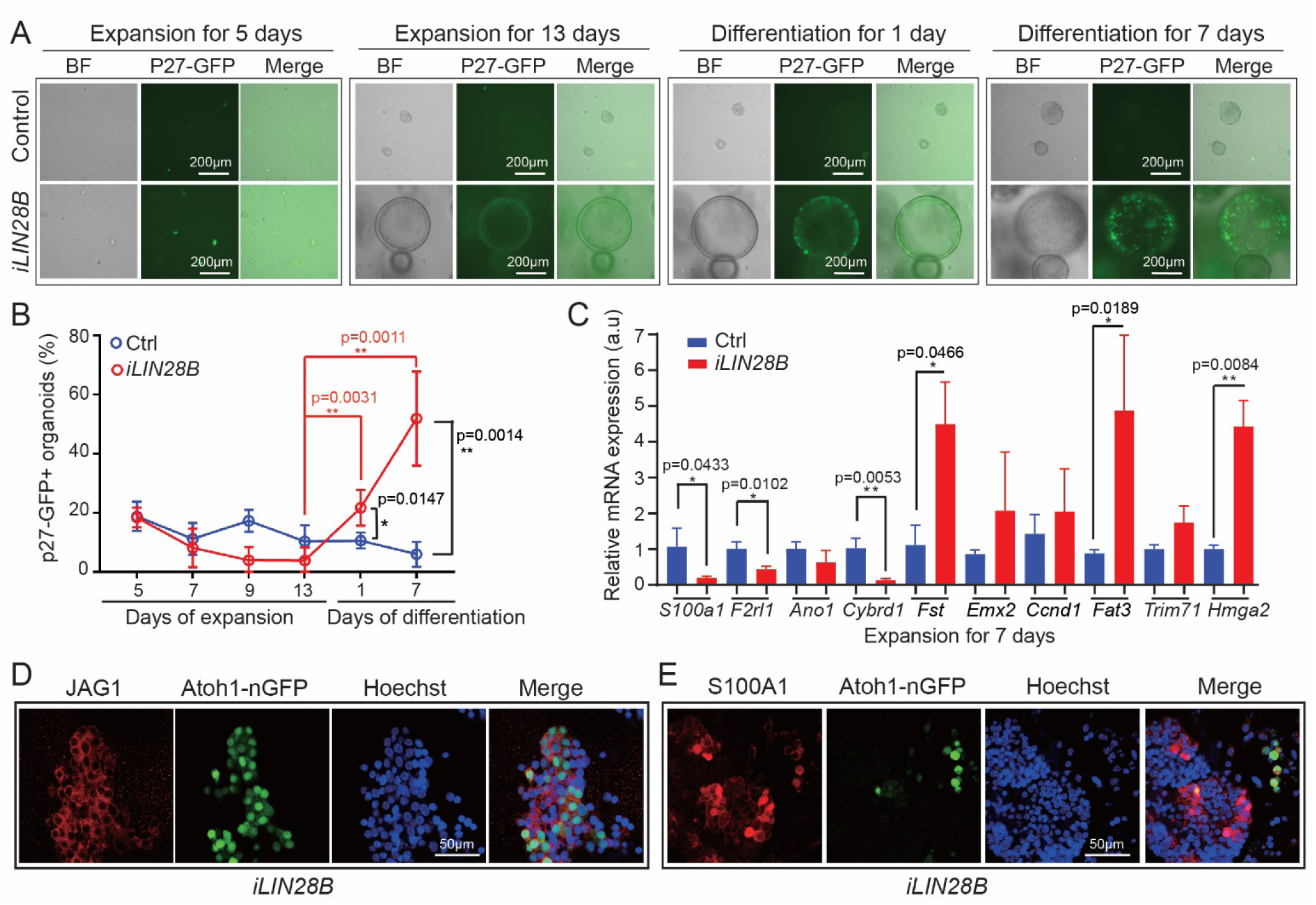
*LIN28B* overexpression promotes supporting cell de-differentiation. Cochlear organoid cultures were established from stage P5 *iLIN28B* transgenic pups and control littermates that lacked the *LIN28B* transgene. Cochlear organoid cultures were maintained as outlined in Fig. 2A. P27-GFP transgene expression was used to monitor post-mitotic supporting cells in *(A-B)*. Atoh1-nGFP transgene expression was used to identify nascent hair cells in (*D and E*). 2-tailed, unpaired Student’s t-test was used to calculate p-values. (*A*) Representative BF and green fluorescent images of p27-GFP expression in LIN28B overexpressing and control organoid cultures at 5 and 13 days of expansion and 1 and 7 days of differentiation. (*B*) Percentage of p27-GFP^+^ organoids in (*A*). Note the percentage of p27-GFP^+^ organoids dramatically increased during differentiation in LIN28B overexpressing (red) cultures (mean ± SD, n=7 animals per group, from 2 independent experiments). (*C*) RT-PCR-based analysis of supporting cell-specific (*S100a1, F2rl1, Ano1, Cybrd1*), pro-sensory cell-specific *(Fst, Fat3, Hmga2, Trim71)* transcripts in control (Ctrl, blue bar) and LIN28B overexpressing (*iLIN28B*, red bar) organoids at 7 days of expansion. *Emx2, a* broad progenitor-specific transcript and cyclin D1 (*Ccnd1*), a pro-sensory/supporting cell-enriched gene, served as controls (mean± SD, n=3 animals per group, from 1 representative experiment, 3 independent experiments). *(D and E)* LIN28B overexpressing organoids after 7 days of differentiation contain Atoh1-nGFP^+^ hair cells (*D* and *E*, Atoh1-nGFP, green) and Atoh1-nGFP^−^ JAGGED1^+^ (*D*, JAG1, red) and Atoh1-nGFP^−^ S100A1^+^ (*E*, S100A1, red) supporting cells. Note that Atoh1-nGFP^+^ S100A1^+^ represent inner hair cells.

To address whether LIN28B overexpression facilitates the de-differentiation of supporting cells-into pro-sensory-like cells, we analyzed the expression of supporting cell (*S100a1, F2rl1, Cybrd1, Ano1*) (33, 48, 49) and pro-sensory cell-specific genes (*Trim71, Hmga2, Fst, Fat3*,)(28, 50) (*Fat3*; Allen Developing Mouse Brain Atlas) in LIN28B overexpressing and control organoids after 7 days of expansion. As controls we analyzed the expression of *Emx2*, which is broadly expressed in cochlear epithelial cells (51), and cyclin D1 *(Ccnd1)*, a pro-sensory-specific gene that continues to be expressed in a subset of supporting cells and hair cells (52). The expression of *Emx2* and cyclin D1 was not significantly changed in LIN28B overexpressing organoids compared to control. However, consistent with supporting cell-de-differentiation, we found that the transcripts of pro-sensory cell-specific genes *Fst, Fat3* and *Hmga2* were 4-fold higher expressed in LIN28B overexpressing organoids compared to control, whereas supporting cell-specific transcripts for *S100a1, F2rl1, Cybrd1* were 2 to 7-fold lower expressed compared to control (Fig. 3 *C*). Once differentiation was induced, LIN28B overexpressing organoids contained both Atoh1-nGFP^+^ hair cells and Atoh1-nGFP^−^ S100A1^+^ or Atoh1-nGFP^−^ JAGGED1^+^ supporting cell-like cell (Fig. S2 *A*)(Fig. 3 *D* and *E*). Together, these results indicate that LIN28B stimulates supporting cells to de-differentiate into pro-sensory-like cells that are capable of producing both hair cells and supporting cells in organoid culture.

### *Let-7* overexpression or loss of LIN28A/B accelerates the age-dependent decline in supporting cell plasticity

LIN28 proteins and *let-7* miRNAs are mutual antagonists. To determine whether higher than normal *let-7* miRNA levels would diminish the regenerative capacity of stage P2 supporting cells, we made use of *iLet-7g* transgenic mice (39). In this double transgenic mouse model, a LIN28A/B resistant form of *let-7g* is expressed under the control of a TRE promoter (*Col1a-TRE-let-7S21L)*, which in the presence of ubiquitously expressed *R26-rtTA-M2* transgene and dox, allows for robust *let-7g* overexpression (28, 39). First, we analyzed whether *let-7g* overexpression disrupts the capacity of young, immature supporting cells to regenerate hair cells in response to Notch inhibition in cochlear explant culture. To ablate hair cells, control and *let-7g* overexpressing cochlear explants were treated with gentamicin for 20 hours, after which Notch inhibitor (LY411575) was added to induce supporting cell-to-hair cell conversion (Fig. S3 *A*). We found that *let-7g* overexpression reduced the production of new hair cell (MYO7A^+^ SOX2^+^) in response to Notch inhibition by more than 2-fold in both intact (PBS, *iLet-7*) and hair cell damaged (gentamicin, *iLet-7*) cochlear explants (Fig. S3, *B* and *C*).

Next, we analyzed whether *let-7g* overexpression inhibits the capacity of young, immature supporting cells to re-enter the cell cycle and propagate in organoid culture. To do so we expanded cochlear epithelia cells from stage P2 *iLet-7g* transgenic animals and their non-transgenic control littermates (lack *let-7g* transgene) in the presence of dox for 10 days (see Fig.1 *A*). We found that overexpression of *let-7g* decreased the number of organoids (Fig. 4 *A* and *C*) and decreased their average size by more than 2-fold (Fig. 4 *A* and *B*). These defects were accompanied by a 30% reduced rate of cell proliferation at 10 days of expansion (Fig. 4 *D* and *E*), as well as defects in supporting cell de-differentiation, as indicated by nearly 2-fold higher *S100a1* expression and more than 3-fold lower *Hmga2* expression in *let-7g* overexpressing organoids compared to control organoids (Fig. 4 *F*). Next, we analyzed the hair cell-producing capacity of control and *let-7g* overexpressing organoids. To obtain sufficient cells for further analysis, we modified the experimental approach and omitted dox from the early phase of expansion. After 5 days of differentiation about 20% of control organoids contained Atoh1-nGFP^+^ cell clusters, whereas only approximately 10% of *let-7g* overexpressing organoids contained Atoh1-nGFP^+^ cell clusters (Fig. 4 *G* and *H*). Moreover, after 5 days of differentiation *iLet-7g* transgenic organoids expressed hair cell-specific genes (*Atoh1, Myo7a*) at 2-fold lower level than control organoids (Fig. 4 *I*). Together, these findings indicate that *let-7* miRNAs antagonize supporting cells proliferating and limit the ability of supporting cells to acquire a hair cell fate. These findings complement our previous finding that *let-7g* overexpression inhibits progenitor cell proliferation during development and inhibits trans-differentiation of supporting cells into hair cells in response to Notch inhibition in cochlear explant culture (28).

**Fig.4.**
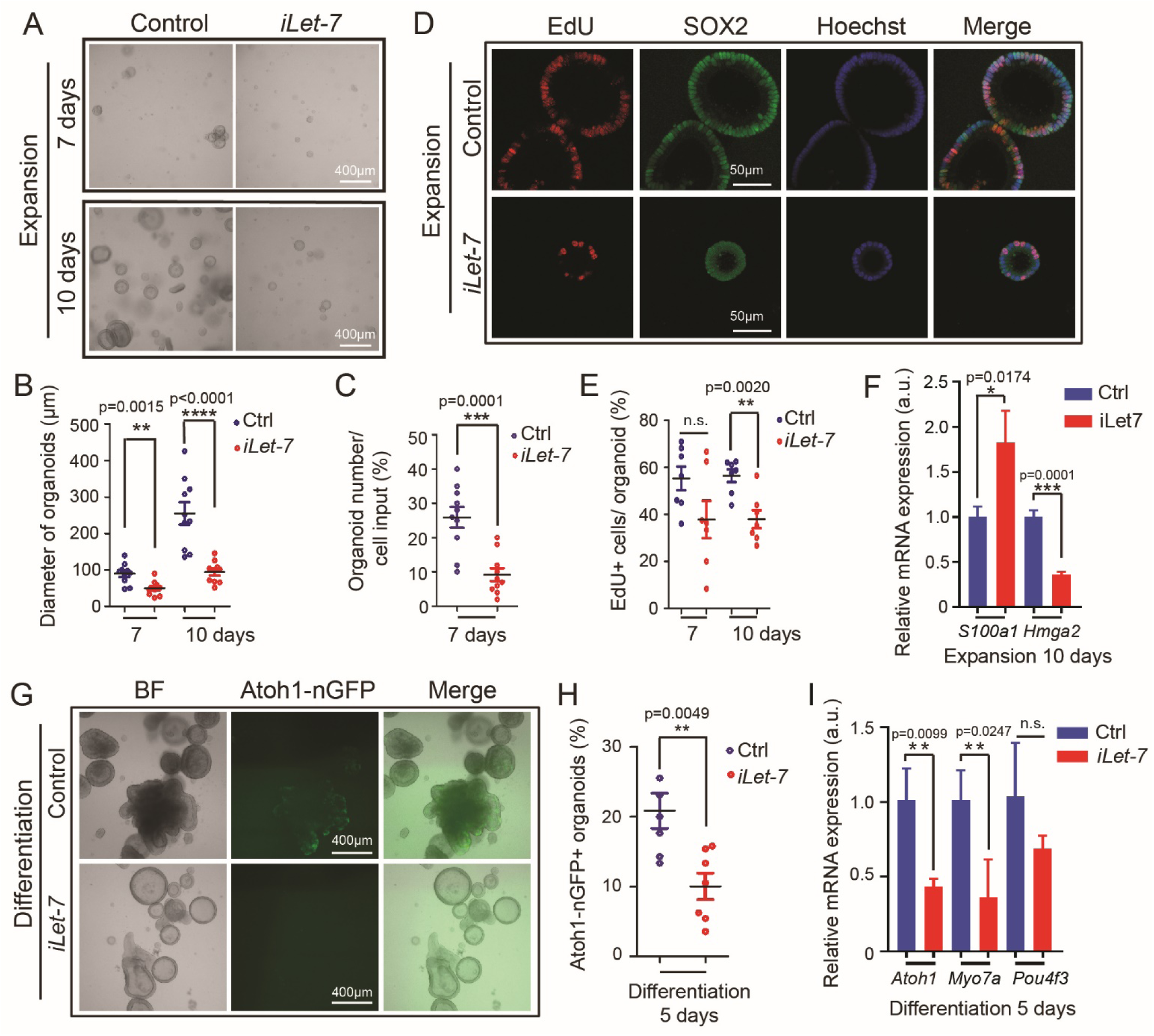
*Let-7g* overexpression inhibits proliferation and hair cell formation in cochlear organoid culture. Cochlear organoid cultures from stage P2 *Atoh1-nGFP; iLet-7* transgenic pups and control littermates that lacked the *let-7g* transgene were maintained as outlined in (Fig. 1 *A*). Bars in *(F)* and *(I)* represent mean ± SD, otherwise individual data points and their mean ± SD were plotted. 2-tailed, unpaired Student’s t-test was used to calculate p-values. (*A*-*F*) Dox was added at plating and dox-containing expansion media was replenished every other day. (*G-I*) Dox was added after 4 days of expansion and was replenished every other day for the remaining 6 days of expansion and 5 days of differentiation. (*A-E*) *Let-7g* overexpression inhibits organoid formation and organoid expansion. (*A*) Bright field (BF) images of control and *let-7g* overexpressing organoid cultures after 7 and 10 days of expansion. (*B*) Diameter of control and *let-7g* overexpressing organoids in (*A*) (n=10 animals per group, from 3 independent experiments). (*C*) Organoid forming efficiency of control and *let-7g* overexpressing organoids (n=10 animals per group, from 3 independent experiments). (*D*) Cell proliferation in control and *let-7g* overexpressing organoids. A single EdU pulse was given at 7 or 10 days (shown) of expansion and EdU incorporation (red) was analyzed 1 hour later. SOX2 (green) marked pro-sensory cells/supporting cells, Hoechst (blue) staining marked cell nuclei. (*E*) Percentage of EdU^+^ cells in (*D*) (n=7 animals per group, from 2 independent experiments). (*F*) RT-PCR of *S100a1* and *Hmga2* mRNA expression in control and *iLet-7* transgenic organoid culture at 10 days of expansion (n=3 animals per group, from 1 representative experiment, 2 independent experiments). (*G*-*I*) *Let-7g* overexpression inhibits hair cell formation. *(G)* BF and green fluorescent images (Atoh1-nGFP) of control and *let-7g* overexpressing organoid cultures. (*H*) Percentage of Atoh1-nGFP^+^ organoids in (*G*) (n=6 animals in control group, n=7 animals in *iLet-7* group, from 2 independent experiments). (*I*) RT-PCR of hair cell-specific (*Atoh1, Myo7a, Pou4f3*) transcripts in control and *let-7g* overexpressing organoids after 5 days of differentiation (n=3 animals per group, from 1 representative experiment, 2 independent experiments). Note that the individual data points in *B*, and *E* represent the average values per animal.

Next, we addressed whether endogenous LIN28A/B protein limits the capacity of neonatal supporting cells to form new hair cells *in vitro*. Mice with single deletion of *Lin28a* or *Lin28b* are viable and show relative mild, or no overt defects, whereas co-deletion of *Lin28a* and *Lin28b* result in severe abnormalities and is embryonic lethal (53). To avoid potential functional redundancy between LIN28B and LIN28A, we conditionally knocked out both *Lin28a* and *Lin28b in vitro* using previously established *Lin28a* and *Lin28b* floxed (f) mice (53) and transgenic mice that ubiquitously express a tamoxifen-inducible form of Cre recombinase (*UBC-CreERT2*) (54). First, we examined, whether endogenous LIN28A/B protein is required for the direct conversion (trans-differentiation) of supporting cells into hair cells (Fig. 5 *A*). We generated cochlear explant cultures from stage P2 *UBC-CreERT2*; *Lin28a*^*f/f*^; *Lin28b*^*f/f*^ pups and littermates that lacked *UBC-CreERT2* transgene (*Lin28a*^*f/f*^; *Lin28b*^*f/f*^). To induce the deletion of *Lin28a* and *Lin28b* (*Lin28a/b*), cultures received 4-hydroxy-tamoxifen (TM) or vehicle control (DMSO) at plating. The next day, cultures were treated with the Notch inhibitor (LY411575) to induce supporting cell-to-hair cell conversion. Three days later, control cochlear explants [(DMSO: *UBC-CreERT2*; *Lin28a*^*f/f*^; *Lin28b*^*f/f*^) or (TM: *Lin28a*^*f/f*^; *Lin28b*^*f/f*^) or (DMSO: *Lin28a*^*f/f*^; *Lin28b*^*f/f*^)] and *Lin28a/b* double knockout (dKO) cochlear explants (TM: *UBC-CreERT2*; *Lin28a*^*f/f*^; *Lin28b*^*f/f*^) were analyzed for newly formed hair cells (MYO7A^+^ SOX2^+^). Consistent with previous reports, we found that supporting cells in control cochlear explant cultures readily converted into new hair cells (MYO7A^+^ SOX2^+^) in response to Notch inhibition, forming close to 20 new hair cells/ 100 µm (Fig. 5 *B* and *C*, blue, red, green). By contrast, *Lin28a/b* dKO cochlear explants produced less than 7 new hair cells/ 100 µm in response to Notch inhibition (Fig 5 B and C, magenta).

**Fig.5.**
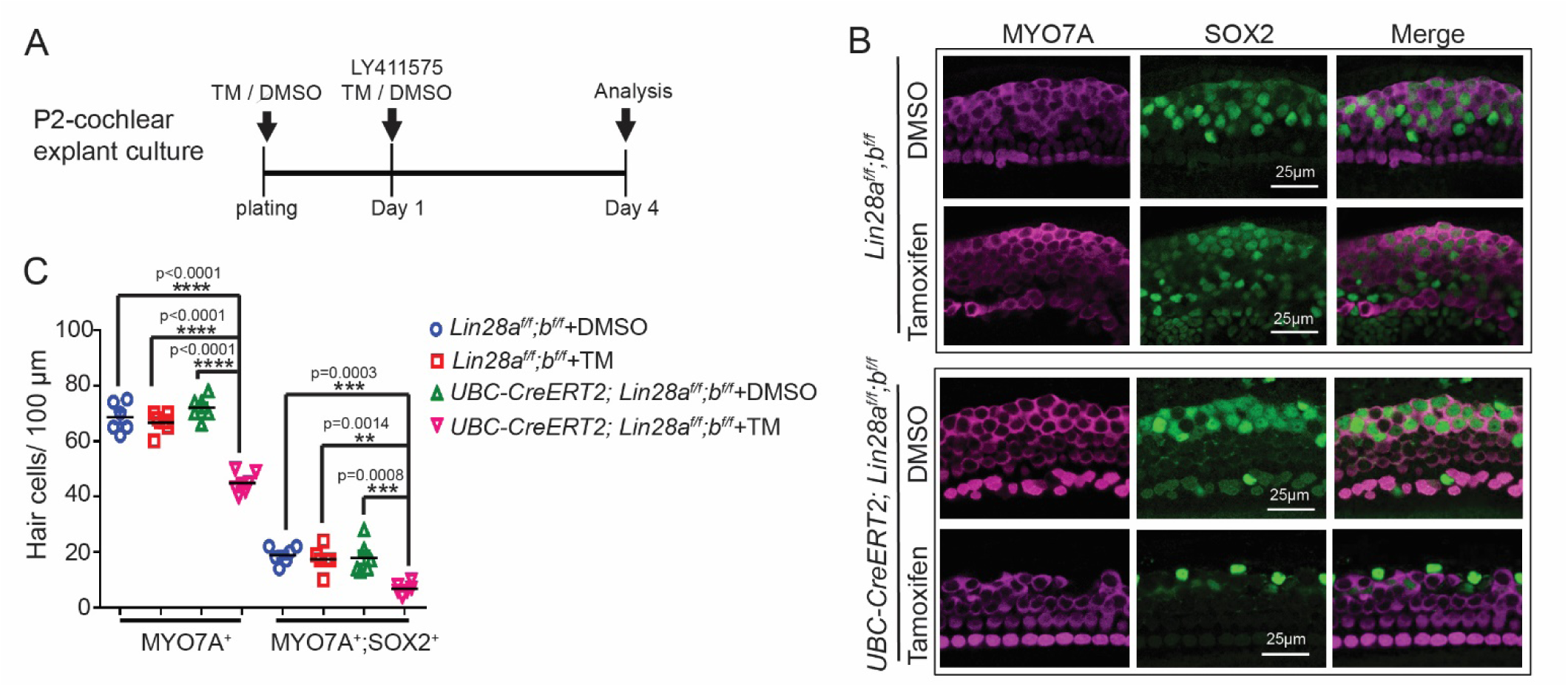
Loss of LIN28A/B inhibits supporting cell-to-hair cell conversion in response to Notch inhibition. (*A*) Experimental strategy. Cochlear explants from stage P2 *UBC-CreERT2; Lin28a*^*f/f*^; *Lin28b*^f/f^ pups and *Lin28a*^*f/f*^; *Lin28b*^*f/f*^ littermates were cultured in the presence of 4-hydroxy-tamoxifen (TM) or DMSO (vehicle control). At day 1 of culture, all cochlear explants received Notch inhibitor LY411575 and 3 days later cochlear explants were analyzed for new hair cells. (*B*) Confocal images of mid-apical turn of control ([TM: *Lin28a*^*f/f*^; *Lin28b*^*f/f*^] or [DMSO: *UBC-CreERT2; Lin28a*^*f/f*^; *Lin28b*^*f/f*^] or [DMSO: *Lin28a*^*f/f*^; *Lin28b*^*f/f*^]) and *Lin28a/b* dKO cochlear explants (TM: *UBC-CreERT2*; *Lin28a*^*f/f*^; *Lin28b*^*f/f*^) immuno-stained for MYO7A (magenta) and SOX2 (green). Note new hair cells expressed MYO7A and SOX2, whereas pre-existing hair cells only expressed MYO7A. (*C*) Quantification of total number of hair cells (MYO7A^+^) as well as newly formed hair cells (MYO7A^+^, SOX2^+^) for control (blue, red, green) and *Lin28a/b* dKO (magenta) in (*B*). Graphed are individual data points, representing average values per animal, and mean ± SD for control (blue, red, green) and *Lin28a/b* dKO (magenta), n=6 mice per group, from 2 independent experiments. 2-way ANOVA with Tukey’s correction was used to calculate p-values.

Next, we determined whether endogenous LIN28A/B is required for young immature supporting cells to propagate and produce new hair cells in cochlear organoid culture (see Fig. 1 *A*). To do so, we established organoid cultures from stage P2 *UBC-CreERT2*; *Lin28a*^*f/f*^; *Lin28b*^*f/f*^ pups and *Lin28a*^*f/f*^; *Lin28b*^*f/f*^ littermates in the presence of 4-hydroxy-tamoxifen (TM) or vehicle control (DMSO). The tamoxifen-induced *Lin28a* and *Lin28b* knock-down was confirmed by RT-qPCR analysis, which showed that *Lin28a/b* dKO organoids expressed *Lin28b* and *Lin28a* transcripts at 3-5-fold lower level than control organoids (Fig. 6 *D*). The reduction in *Lin28a* and *Lin28b* mRNA expression was accompanied by the mild, but a significant down-regulation of the *let-7* target and pro-sensory-specific gene *Hmga2*. Moreover, we found that 2-fold fewer organoids formed in *Lin28a/b* dKO organoid cultures compared to control cultures (Fig. 6 *A* and *C*) and *Lin28a/b dKO* organoids were on average 30% smaller than control organoids (Fig. 6 *A* and *B*). Furthermore, EdU pulse experiments revealed that the fraction of cells that were actively cycling within *Lin28a/b* dKO organoid was ∼30% lower than in control organoids (Fig. 6 *E* and *F*). To address whether LIN28A/B deficiency limits the ability of supporting cells/pro-sensory cells to generate hair cells, we differentiated control and *Lin28a/b* mutant organoids for 10 days and analyzed their expression of hair cell-specific genes (*Atoh1, Pou4f3, Myo7a*) using RT-qPCR. We found that *Lin28a/b* dKO organoids expressed hair cell-specific genes at a more than 3-fold lower level than control organoids (Fig. 6 *G*). Together, our findings indicate that LIN28A/B play an essential role in the mitotic and non-mitotic production of cochlear hair cells *in vitro*.

**Fig.6.**
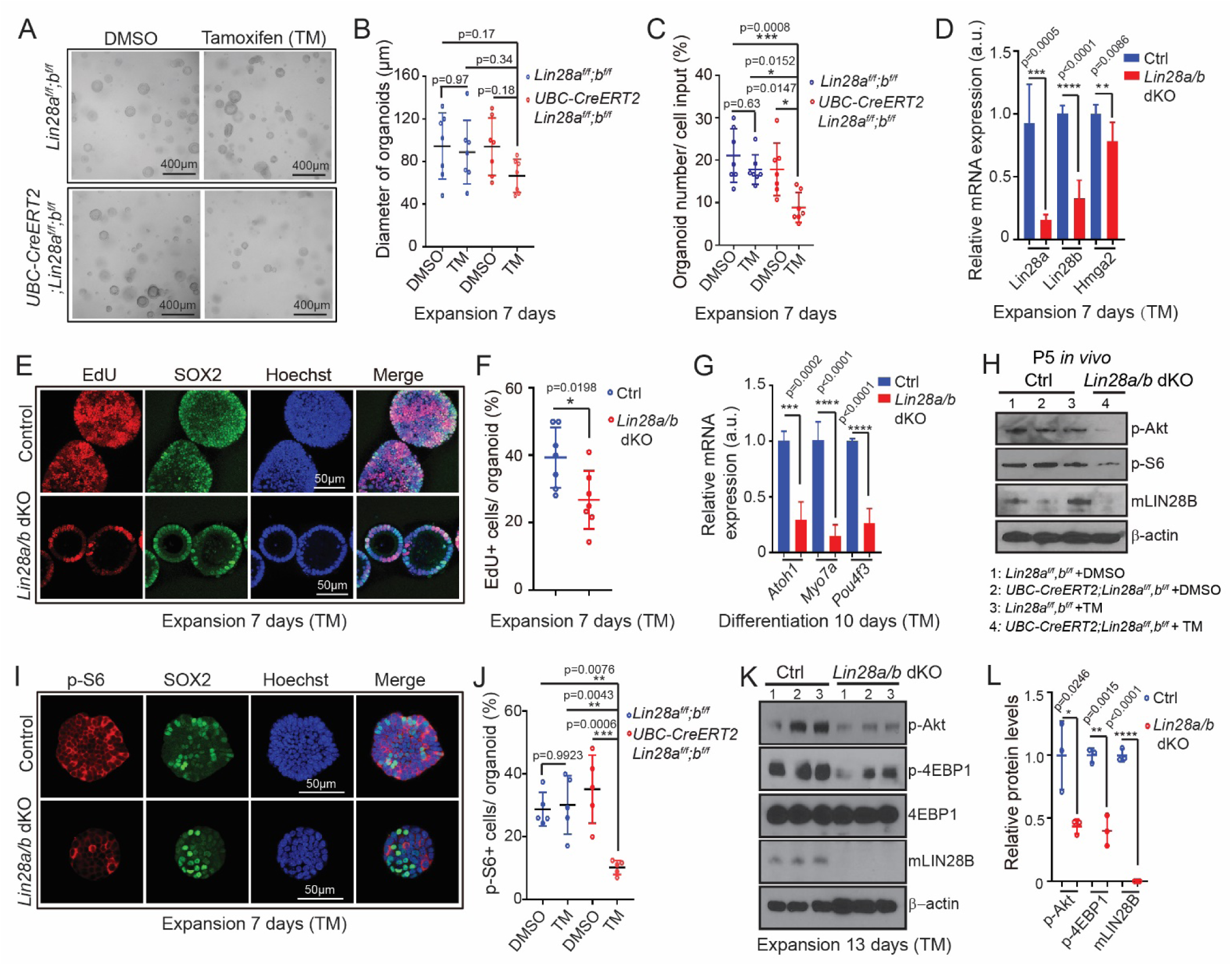
Loss of LIN28A/B attenuates mTOR signaling and limits supporting cell proliferation and hair cell formation in cochlear organoid culture. Cochlear organoid cultures were established from *UBC-CreERT2; Lin28a*^*f/f*^; *Lin28b*^*f/f*^ mice and *Lin28a*^*f/f*^; *Lin28b*^*f/f*^ littermates’ stage P2. Cultures received 4-hydroxytamoxifen (TM) or vehicle control DMSO at plating. SOX2 (green) marks supporting cells/ pro-sensory cells, Hoechst (blue) staining marks cell nuclei. Bars in *(D)* and *(G)* represent mean± SD, otherwise individual data points and their mean ± SD were plotted. *(A-G)* Loss of *Lin28a/b* inhibits cell proliferation and hair cell production in organoid culture. (*A*) Representative BF images of control organoids and *Lin28a/b* dKO organoids at 7 days of expansion. (*B*) Diameter of control and *Lin28a/b* dKO organoids in (*A*) (n=7 animals per group, from 2 independent experiments). (*C*) Organoid forming efficiency in control and *Lin28a/b* dKO cultures (n=7 animals per group, from 2 independent experiments). (*D*) RT-qPCR analysis of *Lin28a, Lin28b* and *Hmga2* mRNA expression in *Lin28a/b* dKO organoids (red bar) compared to control organoids (blue bar) at 7 days of expansion (n=5-6 animals per group, from 2 independent experiments). (*E*) Cell proliferation in control and *Lin28a/b* dKO organoids. A single EdU pulse was given at 7 days of expansion and EdU incorporation (red) was analyzed 1 hour later. (*F*) EdU incorporation in (*E*) (n=7 animals per group, from 2 independent experiments). (*G*) RT-qPCR analysis of *Atoh1, Myo7a* and *Pou4f3* mRNA expression in *Lin28a/b* dKO organoids (red bar) compared to control organoids (blue bar) at 10 days of differentiation (n=4 animals per group, from 2 independent experiments). *(H)* Immunoblots for LIN28B, p-Akt, p-S6 and β-actin using protein lysates of acutely isolated control and Lin28a/b dKO cochlear epithelia, stage P5. (*I*-*L*) Loss of *Lin28a/b* attenuates mTOR signaling in cochlear organoids. (*I*) Immunostaining for p-S6 protein (red) in control and *Lin28a/b* dKO organoids at 7 days of expansion. (*J*) Percentage of p-S6^+^ cells in *(I)* (n=5 animals per group, from 2 independent experiments). (*K*) Immunoblots for p-Akt, p-4EBP1, 4EBP1, mLIN28B and β-actin (loading control) using protein lysates of control and *Lin28a/b* dKO organoids after 7 days of expansion. (*L*) Normalized p-Akt, p-4EBP1 and m-LIN28B protein levels in control and *Lin28a/b* dKO organoids in (*K*) (n=3 animals per group, from 1 representative experiment, 2 independent experiments). 2-way ANOVA with Tukey’s correction was used to calculate p-values in (*B*), (*C*) and (*J*). 2-tailed, unpaired Student’s t-test was used to calculate p-values in (*D*), (*F*), (*G*) and (*L*). Note that the individual data points in (*B*), (*F*) and (*J*) represent the average values per animal.

Recent studies uncovered an important link between LIN28A/B and “The mammalian target of rapamycin” (mTOR) signaling pathway (39, 53). Integrating various growth factor-dependent signals and downstream phosphoinositide 3-kinase (PI3K)-Akt signaling, mTOR signaling pathway promotes anabolic processes in aid of cell growth and cell survival (reviewed in (55)). Central to the mTOR pathway is the serine-threonine kinase mTOR. The mTOR kinase exists in two distinct protein complexes: mTORC1 and mTORC2, which differ in terms of rapamycin sensitivity, substrate specificity and functional outputs (reviewed in ((56)). The rapamycin sensitive mTORC1 complex plays, amongst others, a central role in protein synthesis. Commonly used indicators for mTORC1 activity are the phosphorylated forms of the ribosomal protein S6 (Ser240/244) and the eukaryotic initiation factor 4E binding protein 1 (4E-BP1) (Thr37/46) (57-59). Phosphorylation (p) of the kinase Akt on serine 473 (Ser473) by mTORC2 is a readout for maximal Akt activation downstream of PI3K (60).

To determine whether similar to LIN28B, mTOR activity declines during cochlear maturation, we analyzed p-S6 and p-Akt protein levels in cochlear epithelial lysates from stage P2 and P5 wild type mice. We found that similar to the drop in LIN28B protein, p-S6 and p-Akt protein levels were significantly reduced in P5 cochlear sensory epithelia compared to P2 (Fig. S1 *B* and *C*). Furthermore, similar to endogenous LIN28B, mTOR activity (p-4E-BP1) was higher in the apical portion of stage P5 cochlear sensory epithelia than the basal one. However due the high variability across samples the apex-base difference in mTOR activity was not statistical significant (Fig. S1 *D-F*). To determine whether LIN28A/B positively regulates mTOR activity in cochlear epithelial cells *in vivo*, we analyzed mTOR activity in acutely isolated cochlear epithelial lysates after postnatal LIN28B overexpression (*iLIN28B)* or LIN28A/B knockdown (*Lin28a/b* dKO, *iLet-7g)*. We found that conditional ablation of *Lin28a/b* (Fig. 6 H) or *let-7g* overexpression in neonatal mice (Fig. S4 *A-C*) reduced cochlear m-TOR activity (p-Akt and or p-S6 protein), whereas LIN28B overexpression increased it (Fig. S5 *A and B*).

Consistent with these *in vivo* findings, p-Akt, p-S6 and or p-4E-BP1 protein levels were 1.5 to 2-fold lower in lysates of LIN28A/B deficient organoids (*Lin28a/b* dKO, *iLet-7g*) than control organoids (Fig. 6 *K* and *L*) (Fig. S4 *F* and *G*). Furthermore, the percentage of p-S6^+^ cells in LIN28A/B deficient organoids (*Lin28a/b dKO, iLet-7g*) was 2 to 3-fold lower than in control organoids (Fig. 6 *I and J*)(Fig. S4 *D* and *E*). Conversely, the levels of p-Akt and p-S6 proteins were more than 1.7-fold higher in lysates of LIN28B overexpressing organoids compared to control organoid lysates (Fig. S5 *C* and *D*). Moreover, we found that LIN28B overexpression increased the percentage of p-S6^+^ cells within organoids by more than 2-fold (Fig. S5 *E and F*). Co-staining with SOX2 revealed that the majority of p-S6^+^ cells in LIN28B overexpressing organoids expressed SOX2 at a low level (Fig. S5 *G*; SOX2^low^, p-S6^high^; red dots).

Next we determined whether the expansion of supporting cells/pro-sensory cells requires mTOR signaling. To block mTOR signaling in organoid culture we used the mTORC1 complex inhibitor rapamycin (reviewed in (61)). 50 µM of rapamycin has been reported to induce hair cell death rat cochlear explants (62). However, pilot experiments revealed that rapamycin at 4 ng/mL (4.4 µM) effectively blocked mTOR activity (Fig. S6 *A* and *B*), and attenuated the growth of P2 wild type cochlear organoids (Fig. S6 *C-E*), without increasing cell death within these organoids (Fig. S6 *F* and *G*). By contrast, 50 ng/mL (56 µM) rapamycin did induce cell death in organoid culture, consistent previous reports (Fig. S6 *H*). Furthermore, we found that the presence of 4 ng/mL of rapamycin reversed the LIN28B-mediated increase in organoid forming efficiency and organoid growth in P5 organoid culture (Fig. 7 *A*-*C*). Next, we investigated whether mTORC1 activity is required for the formation of hair cells in P5 LIN28B overexpressing organoid cultures. To avoid the negative effects of rapamycin on organoid formation and expansion, we added rapamycin or vehicle control (DMSO) to LIN28B overexpressing organoids just prior to the first formation of Atoh1-nGFP+ cells at day 10 of expansion. The initial induction of Atoh1-nGFP^+^ cells within organoids occurred in rapamycin treated cultures at a similar rate than in control cultures (Fig. 7 *D* and *E*). However, LIN28B overexpressing organoids treated with rapamycin contained 3.5-fold fewer Atoh1-nGFP^+^ MYO7A^+^ hair cells and expressed hair cell-specific genes (*Myo7a, Pou4f3*) at a 2-fold lower level than LIN28B overexpressing organoids treated with vehicle control (Fig. 7 *F*-*H*). To address whether mTOR signaling plays a role in the initial induction of *Atoh1*, we modified our experimental approach and pre-treated the culture media with rapamycin 1 day prior to the induction of LIN28B overexpression at day 8 of expansion (Fig. S7 *A*). Using this modified experimental approach, we found that rapamycin treatment significantly reduced the number of Atoh1-nGFP+ organoids produced in response to LIN28B overexpression (Fig. S7 *B* and *C*).

**Fig.7.**
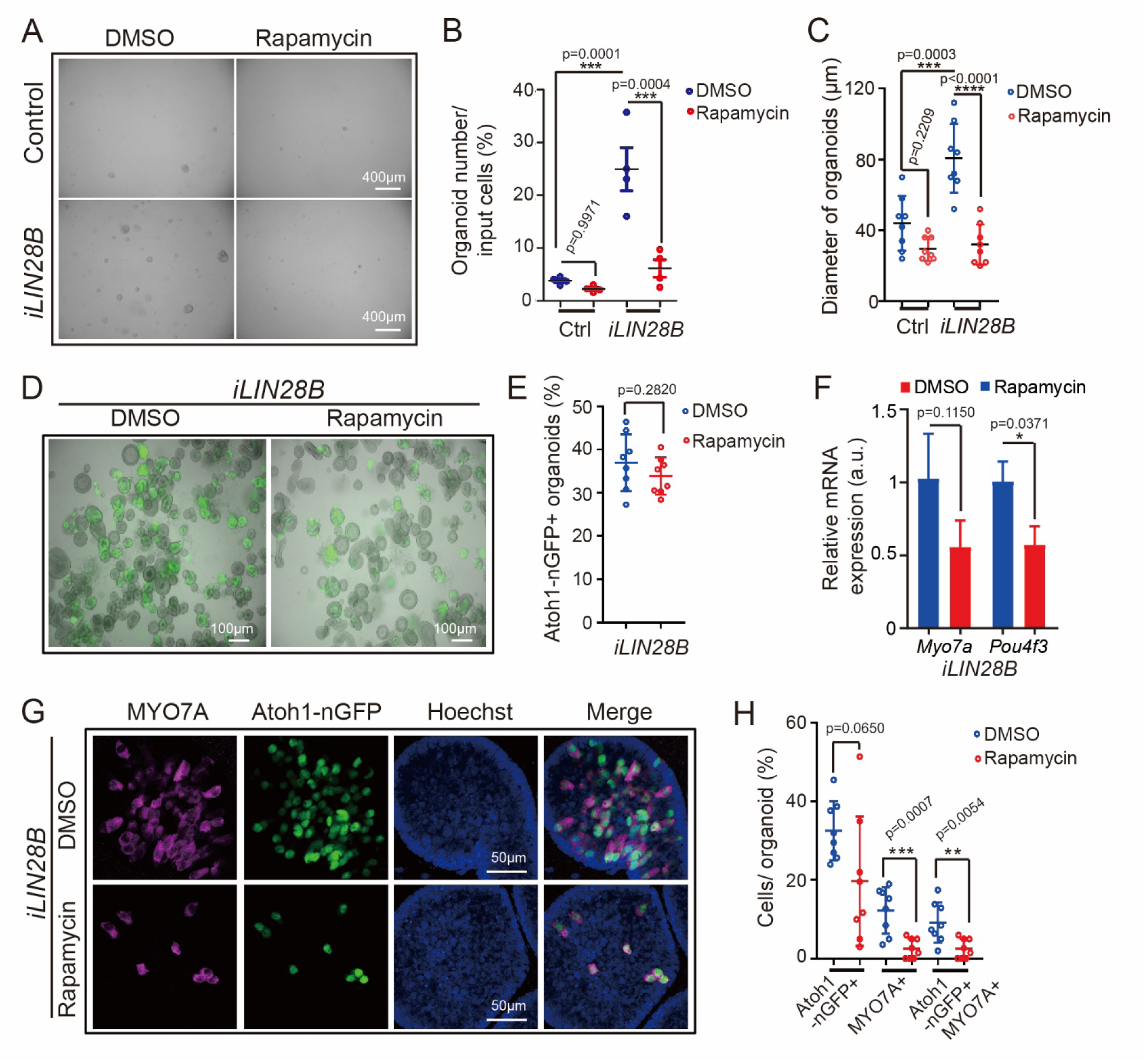
Inhibition of mTOR signaling attenuates LIN28B-induced organoid growth and hair cell formation. Cochlear organoid cultures were established from stage P5 *Atoh1-nGFP*; *iLIN28B* transgenic mice and *Atoh1-nGFP* transgenic control littermates and maintained as outlined in (Fig. 2 *A*). Dox-containing culture media was replenished every other day. Graphs in *(B), (C), (E)* and *(H)* show individual data points and the mean ± SD. (*A*-*C*) Rapamycin attenuates LIN28B’s positive effect on cell growth in organoid culture. Rapamycin (4 ng/ml) or vehicle control (DMSO) was added at plating and replenished every other day. (*A*) BF images of control and LIN28B overexpressing (*iLIN28B*) organoids after expansion in the presence of DMSO or rapamycin for 7 days. (*B*) Organoid forming efficiency in (*A*) (n=4 animals per group, from 2 independent experiments). (*C*) Organoid diameter in (*A*) (n=8 animals per group, from 3 independent experiments). (*D*-*H*) Rapamycin attenuates LIN28B’s positive effect on hair cell formation in organoid culture. Rapamycin (4 ng/ml) or DMSO was present during the final phase of expansion (10-13 days) and during the 5 days of differentiation. (*D*) Merged BF and green fluorescent images (Atoh1-nGFP) of LIN28B overexpressing organoid cultures treated with DMSO or rapamycin. (*E*) Percentage of Atoh1-nGFP^+^ organoids in (*D*) (n=8 animals per group, from 2 independent experiments). (*F*) RT-qPCR-based analysis of *Myo7a* and *Pou4f3* transcripts in LIN28B overexpressing organoids treated with rapamycin (red bar) or DMSO (blue bar) (mean ± SD, n=2 animals per group, from 1 representative experiments, 2 independent experiments total). (*G*) Confocal images LIN28B overexpressing organoids cultured in the presence of rapamycin or DMSO. Newly formed hair cells are identified by their co-expression of Atoh1-nGFP (green) and MYO7A (magenta). Nuclei were counterstained with Hoechst (blue). (*H*) Percentage of GFP^+^ (Atoh1-nGFP), MYO7A^+^ and GFP^+^ MYO7A^+^ hair cells per organoid (*F*) (n=8 animals per group, from 2 independent experiments). 2-tailed, unpaired Student’s t-test was used to calculate p-values in (*E*), *(F)* and (*H*). 2-way ANOVA with Tukey’s correction was used to calculate p-values in (*B*) and (*C*). Note that the individual data points in (*C*), *(E)* and (*H*) represent average values per animal.

Next, we analyzed whether elevated LIN28B-mTOR levels would promote the production of new hair cells in stage P5 cochlear explants. To ensure robust *LIN28B* transgene expression, we administered dox *in vivo* starting at stage E17.5 and provided dox throughout the duration of the experiment. At plating, stage P5 control and LIN28B overexpressing cochlear explants received rapamycin or DMSO (vehicle control). The next day the Notch inhibitor LY411575 and Wnt-activator (GSK3-β inhibitor) CHIR99021 was added to the culture media and 3 days later explants were harvested and analyzed for newly generated hair cells (Atoh1-nGFP^+^ SOX2^+^ MYO7A^+^) (Fig. 8 *A*). Inhibition of Notch signaling in combination with Wnt-activation is highly effective in stimulating hair cell production in the neonatal cochlea *in vitro* and *in vivo* (22, 63). However, the effectiveness of such combined treatment has not yet been tested at later stages. We found that combined Notch inhibition and Wnt activation yielded on average 10 new hair cells/100 µm in control cochlear explants, a modest but significant improvement over the previously reported failure to produce hair cells with Notch inhibitor treatment alone (26) (Fig. 8 *B* and *C*, control and DMSO). As anticipated, LIN28B overexpression significantly increased the production of new hair cells in P5 cochlear explants, yielding on average 25 new hair cells/100 µm, a more than 2-fold improvement compared to control cochlear explants (Fig. 8 *B* and *C, iLIN28B* and DMSO). The increase in hair cell formation in response to LIN28B overexpression was not due to an increase in cell proliferation as new hair cells were produced by non-mitotic mechanisms (Fig. 8 *B*, EdU). The ability to produce new hair cells was completely abolished when control cochlear explants were cultured in the presence of rapamycin (Fig. 8 *B* and *C*, control and rapamycin). Furthermore, we found that rapamycin treatment completely reversed the gain in hair cell production observed in response to LIN28B overexpression (Fig. 8 *B* and *C, iLIN28B* and rapamycin).

**Fig.8.**
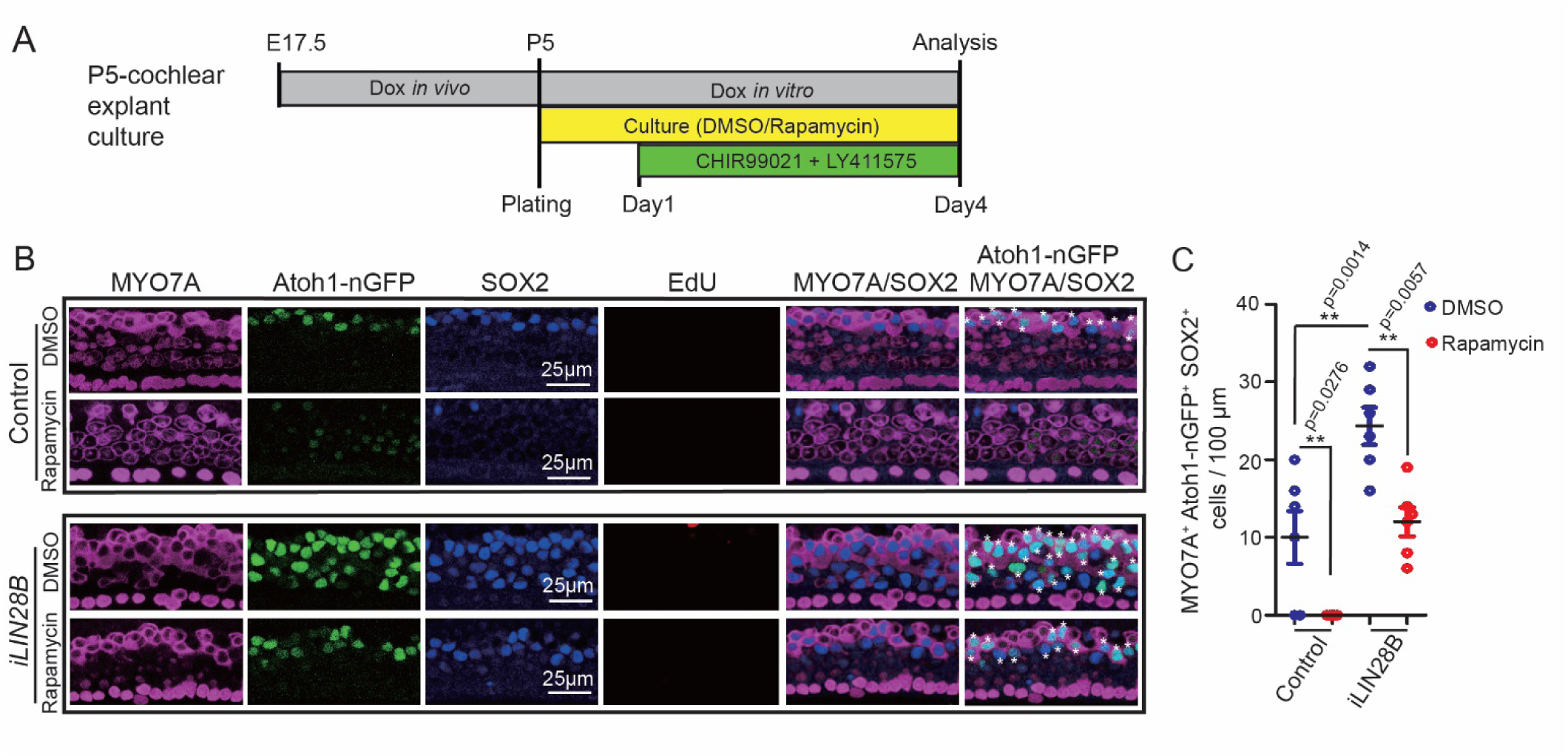
LIN28B promotes non-mitotic hair cell production in an mTOR-dependent manner. (*A*) Experimental strategy. Cochlear explant cultures were established from stage P5 *Atoh1-nGFP iLIN28B* transgenic and *Atoh1-nGFP* transgenic control littermates. Pregnant dams were fed dox containing food starting at E17.5 and dox was present throughout cochlear explant culture. EdU (3 µM), rapamycin (4 ng/mL) or vehicle control DMSO was added at plating. CHIR99021 (3 μM) and LY411575 (5 μM) were added the next day. The culture media was replenished every day. (*B*) Rapamycin treatment inhibits hair cell generation in control and LIN28B overexpressing cochlear explants. Shown are representative confocal images of the mid-apical turn of control and *LIN28B* overexpressing cochlear explants labeled for MYO7A (magenta), Atoh1-nGFP (green), SOX2 (blue) and EdU (red). White asterisks mark newly formed hair cells. (*C*) Quantification of newly formed hair cells (MYO7A^+^; Atoh1-nGFP^+^; SOX2^+^) in (*B*) (graphed are average values for each animal and the mean± SD, n=6 animals per group, from 2 independent experiments, 2-way ANOVA with Tukey’s correction was used to calculate p-values).

## DISCUSSION

The developing embryo and neonate show a remarkable capacity for regeneration in response to injury; however, only few tissues retain such regenerative plasticity into adulthood (reviewed in (64)). Likewise, the ability of murine cochlear supporting cells to produce hair cells sharply declines during the first postnatal week (27, 48) and little to no hair cell production/ regeneration is observed within the cochlea of adult mice (14, 17, 65). The reason for the decline in supporting cell plasticity is unknown.

Here, we provide evidence that the decline in supporting cell plasticity during cochlear maturation is, at least in part, the result of diminishing LIN28B-mTORC1 activity. We show that *in vivo*, coinciding with the abrupt decline in supporting cells plasticity, cochlear LIN28B protein levels and Akt-mTORC1 kinase activity sharply decline. We establish a regulatory link between cochlear Akt-mTORC1 activity and LIN28B/*let-7* expression levels *in vitro* and *in vivo*, with Akt-mTORC1 kinase activity being augmented by LIN28B overexpression but suppressed by loss of LIN28A/B or overexpression of *let-7g*. Using cochlear organoid and explant culture systems, we found that diminished LIN28B-mTORC1 activity, in response to *let-7g* overexpression, or targeted deletion of *Lin28a/b*, accelerated the developmental decline in supporting cell plasticity. Conversely, we found that LIN28B overexpression halted the decline and enabled supporting cells that were undergoing maturation to activate a progenitor-like state and to produce new hair cells in responds to regenerative cues. Finally, using the mTORC1 inhibitor rapamycin, we uncovered that LIN28B-induced supporting cell reprogramming required mTORC1-dependent signaling.

Rapamycin sensitive mTOR signaling supports a wide range of regenerative processes including the regrowth of axons, injury-induced cell proliferation and stem cell-based muscle regeneration (66-69). Activation of mTORC1 kinase activity in response to injury or other environmental stimuli increases the cell’s synthesis of proteins and other macromolecules and ramps up mitochondrial ATP production, which is critical for the cell’s ability to divide and/or differentiate (70). Thus, it is likely that rapamycin sensitive mTOR signaling enhances the regenerative capacity by reprogramming the metabolism of supporting cells, ensuring that supporting cells have the energy and building blocks needed for mitotic or non-mitotic hair cell generation. Indeed, LIN28A/B induced and mTORC1-dependent metabolic reprogramming has been reported in the context of organismal growth and glucose metabolism (39, 53). Moreover, mTOR signaling is known to cross-talk and cross-regulate growth-related signaling pathways including canonical Wnt/β-catenin signaling (71)(reviewed in (72)). Future studies will examine both LIN28B-dependent an LIN28B-independent roles of mTOR signaling in supporting cell plasticity and address how mTOR signaling may alter the strength and transcriptional output of Wnt/β-catenin signaling in the context of hair cell regeneration.

How does LIN28B promote mTORC1 activity in cochlear epithelial cells? Several members of the mTOR pathway and the upstream IGF2/IGF1R and amino acid sensing pathways are direct *let-7* targets (39, 73, 74), suggesting that LIN28B enhances mTOR activity, at least in part, by relieving *let-7* mediated repression. In addition, LIN28A/B have been shown to promote IGF2/IGF1R–mTOR signaling through increasing the mRNA translation of key member genes (39, 75, 76). It needs to be noted that treatment with the mTORC1 inhibitor rapamycin attenuated, but did not completely block, supporting cell proliferation and hair cell production, suggesting that LIN28B promotes these processes through additional, mTORC1 independent mechanisms. LIN28A/B and *let-7* miRNAs promote self-renewal and regeneration via translational activation/ repression of pro-growth genes such as *Igf2bp1* (77), *Trim71* (78) and *Hmga2* (79), as well via the direct targeting of genes critical for oxidative phosphorylation (80) and lipid synthesis (81). Indeed, we found LIN28B activated, whereas *let-7g* limited *Hmga2* expression in cochlear organoids. In addition, we found that LIN28B overexpression led to a transient downregulation in *Sox2* mRNA expression at the peak of organoid expansion. Recent studies found that SOX2 positively regulates *p27/Kip1* transcription and reduction in *Sox2* expression primes cochlear supporting cells for proliferation and hair cell regeneration (47, 65). Future studies are warranted to address LIN28B’s role in *Sox2* expression during hair cell regeneration.

It is important to note that re-activation of LIN28B is, by itself, not sufficient to stimulate supporting cell proliferation and is not sufficient to induce the conversion of supporting cells into hair cell. Additional signals and factors, such as Wnt over-activation and or Notch inhibition are required to stimulate cochlear supporting cell proliferation and their conversion into hair cells. Furthermore, it needs to be emphasized that our study was conducted *in vitro* and was limited to immature, early postnatal stages. While recent studies showed promise in regenerating vestibular hair cells in adult mice (9, 82), the regeneration of cochlear hair cells in adult mice has proven thus far an impossible task (65) and reviewed in (83).

Similar to mammals, auditory supporting cells in birds are highly specialized post-mitotic cells. The lack of progenitor-like features has led to the yet to be proven hypothesis that hair cell loss in birds triggers a partial de-differentiation of supporting cells, in which supporting cells revert to a transitional progenitor-like state (reviewed in (84)). Here, we provide evidence that supporting cells do undergo a process of de-differentiation in organoid culture and show that supporting cell-de-differentiation is regulated by the LIN28B/*let-7* axis. We found that overexpression of *let-7g* in cochlear organoids repressed the downregulation of supporting cell-specific genes (e.g. *S100a1*) and limited the upregulation of pro-sensory-specific genes (*Hmga2)* whereas LIN28B reactivation enhanced it. Such positive role of LIN28B in supporting cell de-differentiation is reminiscent of the role of LIN28A in Müller glia de-differentiation. Prior studies found that in zebrafish *Lin28a* reactivation is essential for the de-differentiation of Müller glia into multi-potent retinal progenitors in response to injury (85, 86). The importance of LIN28-mediated de-differentiation in the context of retinal regeneration is further highlighted by the recent finding that LIN28A reactivation induces Müller glia de-differentiation and proliferation in adult mice (87, 88).

We anticipate that the here presented findings will lead to a similar evaluation of the role of LIN28A/B in injury-induced hair cell regeneration in birds and fish. Furthermore, an evaluation of the role of LIN28A/B in vestibular hair cell regeneration may address why vestibular supporting cells have a higher regenerative capacity than their cochlear counterparts (89). Most importantly, our observation that re-activation of LIN28B facilitates the reprogramming of supporting cells into “progenitor-like” cells, may inform future therapeutic strategies that combine LIN28B-mediated reprograming with Wnt-activation and/or Notch inhibition to regenerate lost cochlear hair cells in adult mammals.

## METHODS

### Mouse breeding and genotyping

All experiments and procedures were approved by the Johns Hopkins University Institutional Animal Care and Use Committees protocol, and all experiments and procedures adhered to National Institutes of Health-approved standards. The *Atoh1/nGFP* transgenic (tg) (MGI:3703598) mice were obtained from Jane Johnson (University of Texas Southwestern Medical Center, Dallas) (37). The *p27/GFP tg* mice (MGI:3774555) were obtained from Neil Segil (University of Southern California, Los Angeles) (44). The *Col1a-TRE-LIN28B* (*MGI:5294612), Col1a-TRE-let-7S21L* (MGI:5294613) and *Lin28a*^*fl/fl*^ (MGI:5294611) and *Lin28b*^*fl/fl*^ (MG:5519037) mice were obtained from George Q. Daley (Children’s Hospital, Boston) (39, 53). *UBC-CreERT2* tg (MGI:3707333) (stock no. 007001) (54) and *R26-M2-rtTA* (MGI:3798943) (stock no. 006965) mice were purchased from Jackson Laboratories (Bar Harbor, ME). Mice were genotyped by PCR and genotyping primers are listed in Table S1. Mice of both sexes were used in this study. Embryonic development was considered as E0.5 on the day a mating plug was observed. To induce *LIN28B* or *let-7g* transgene expression, doxycycline (dox) was delivered to time-mated females via ad libitum access to feed containing 2 grams of doxycycline per kilogram feed (Bioserv, no. F3893).

### Organoid culture

Cochlear tissue from early postnatal mice (postnatal days 2 or 5) was harvested and micro-dissected in Hank’s balanced salt solution (HBSS). Enzymatic digest with dispase (1 mg/mL, Thermo Fisher Scientific, no. 17105041) and collagenase (1 mg/mL, Worthington, no. LS004214) was used to isolate cochlear epithelia as previously described (90). To obtain a single cell suspension, cochlear epithelia were incubated in TrypLE solution (ThermoFisher, no. 2604013), triturated and filtered through a 35 µm filter (BD, no. 352235). In each experiment an aliquot of cells was counted to establish the total number of cells isolated per animal. Equal numbers of cells were then resuspended in a 1:1 mix of Matrigel (Corning, no. 356231) and expansion medium and plated into pre-warmed 4-well plates (CELLTREAT, no. 229103). For mice stage P2, 8 separate cochlear organoid cultures, placed in individual wells were established. For mice stage P5, 5 separate cochlear organoid cultures, placed in individual wells were established. The expansion medium was prepared with DMEM/F12 (Corning, no. 10–092-CV), N-2 supplement (1X, ThermoFisher, no.17502048), B-27 supplement (1X, ThermoFisher, no.12587010), EGF (50 ng/mL, Sigma-Aldrich, no. SRP3196), FGF2 (50 ng/mL, ThermoFisher, no. PHG0264), CHIR99021 (3 μM, Sigma-Aldrich, no. SML1046), VPA (1 mM, Sigma-Aldrich, no. P4543), 616452 (2 μM, Sigma-Aldrich, no. 446859-33-2) and penicillin (100 U/mL, Sigma-Aldrich, no. P3032). Doxycycline hyclate (10 μg/mL, Sigma-Aldrich, no. D9891) and 4-hydroxy-tamoxifen (20 ng/mL, Sigma-Aldrich, no. H7904) was added to cultures on the first day of expansion. To induce differentiation, expansion media was replaced by a differentiation media composed of DMEM/F12, N2 (1X), B27 (1X), CHIR99021 (3 μM) and LY411575 (5 μM, Sigma-Aldrich, no. SML0506). Culture media was changed every other day.

### Quantification of organoid forming efficiency, organoid diameter and GFP reporter expression

Bright field and fluorescent images of organoid cultures were captured with an Axiovert 200 microscope using 5X and 10X objectives (Carl Zeiss Microscopy). To calculate organoid forming efficiency, the total number of organoids per culture was counted manually and values were normalized to the total number of cells plated. To calculate organoid size, the diameter of organoids in two randomly chosen fields was measured using ImageJ (https://imagej.nih.gov/ij/)) and the average value per animal was reported. To establish the percentage of GFP^+^ organoids per culture, the total number of organoids and the number of GFP^+^ organoids was counted manually. For each genotype and treatment, three independent organoid cultures were established and analyzed. At a minimum, two independent experiments were conducted and analyzed.

### Explant culture

Cochlear capsule from individual pup’s stage P2 or P5 were removed in Hank’s Balanced Salt Solution (Corning, no. 21–023-CV) and the remaining tissue, including the cochlear sensory epithelium and the innervating spiral ganglion, was placed onto SPI-Pore membrane filters (Structure Probe, no. E1013-MB) and cultured in DMEM-F12 (Corning, no. 10–092-CV), EGF (5 ng/mL, Sigma-Aldrich, no. SRP3196), penicillin (100 U/mL, Sigma-Aldrich, no. P3032) and N2 Supplement (1X, ThermoFisher, no. 17502048). All cultures were maintained in a 5% CO2/ 20% O2 humidified incubator. To induce *let-7g* or *LIN28B* transgene expression, timed pregnant dams received dox-containing feed starting at E17.5/<u>E18.5</u> until pups were harvested and their cochlear tissue processed for cochlear explant culture. To maintain *let-7g* or *LIN28B* transgene expression *in vitro*, the culture media was supplemented with doxycycline hyclate (10 μg/mL, Sigma-Aldrich, no. D9891). In *iLet-7* experiments, cochlear explants were established from stage P2 dox-induced *iLet-7* pups and control littermates. To induce hair cell loss, cochlear explants received gentamicin sulfate (100µg/mL, Sigma-Aldrich, no. G1272) at plating. After 20 hours the gentamicin-containing media was replaced with culture media containing LY411575 (5 μM, Sigma-Aldrich, no. SML0506). For iLIN28B experiments, cochlear explants were established from stage P5 dox-induced iLIN28B pups and control littermates. To block mTORC1 activity, cochlear explants received rapamycin (4 ng/mL Sigma-Aldrich, no. R0395) or vehicle control DMSO at plating. CHIR99021 (3 μM, Sigma-Aldrich, no. SML1046), LY411575 (5 μM, Sigma-Aldrich, no. SML0506) and EdU (3 μM, Thermo Fisher, no. C10338) were added the following day (day 1). For *Lin28a/b* dKO experiments, *UBC-CreER*; *Lin28a*^*f/f*^; *Lin28b*^*f/f*^ and *Lin28a*^*f/f*^; *Lin28b*^*f/f*^ mice were harvested at stage P2. To induce Cre mediated deletion 4-hydroxy-tamoxifen was added to the culture medium at plating. LY411575 (5 μM) or vehicle control DMSO were added the next day. In all experiments, the culture media were exchanged every other day, and cultures were maintained for a total of 4 days, after which they were processed for immuno-staining.

### Immunohistochemistry

Organoids/explants were fixed with 4% (vol/vol) paraformaldehyde in PBS (Electron Microscopy Sciences, no. 15713) for 30 minutes at room temperature. To permeabilize cells and block unspecific antibody binding, organoids/ explants were incubated with 5% (vol/vol) TritonX-100/ 10% (vol/vol) FBS in 1X PBS for 30 minutes. The following primary antibodies were used: mouse anti-Calretinin (Sigma-Aldrich, no. MAB1568, 1:500), goat anti-JAG1 (Santa Cruz, no. sc-6011, 1:500), rabbit anti-myosin VIIa (Proteus Biosciences, no. 25-6790, 1:500), goat anti-prestin (Santa Cruz, no. sc-22692, 1: 500), rabbit anti-S100A1 (Abcam, no. ab11428, 1:500), goat anti-SOX2 (Santa Cruz Biotechnology, no. sc-17320, 1:500) and rabbit anti-p-S6 (Ser240/244) (Cell Signaling, no. 5364, 1:500). The secondary antibodies used were, donkey anti-rabbit IgG (H+L) Alexa Fluor 546 (1:1000, ThermoFisher, no. A10040), donkey anti-rabbit IgG (H+L) Alexa Fluor 647 (1:1000, ThermoFisher, no. A-31573), donkey anti-mouse IgG (H+L) Alexa Fluor 546 (1:1000, ThermoFisher, no. A-10036), donkey anti-goat IgG (H+L) Alexa Fluor 488 (1:1000, ThermoFisher, no. A-11055) and donkey anti-goat IgG (H+L) Alexa Fluor 546 (1:1000, ThermoFisher, no. A-11056). Antibody labeling was performed according to manufacturer’s recommendations. Cell nuclei were stained using Hoechst 33258 solution (Sigma-Aldrich, no. 94403).

### Cell proliferation

EdU (5-ethynyl-2’-deoxyuridine) was added to culture media at a final concentration of 3 μM. EdU incorporation was detected using Click-iT™ Edu Alexa Fluor™ 555 imaging Kit (ThermoFisher, no. C10338) following the manufacturers specifications.

### Cell death

TUNEL reactions were used to detect dying cells. PFA-fixed and permeabilized organoids were processed using the In Situ Cell Death Detection Kit, Fluorescein (Roche, no.11684795910) according to manufacturer’s specifications. As positive control, organoids were pre-incubate 10 min at room temperature in DNase I (3000 U/ml, in 50 mM Tris-HCl, pH 7.4, 1mg/ml BSA). As negative control, organoids were processed without a reaction mixture that lacked terminal deoxynucleotidyl transferase.

### Cell counts

High power confocal single plane and z-stack images of fluorescently immuno-labeled organoids and explants were taken with 40X objective using LSM 700 confocal microscope (ZEISS Microscopy). To establish the percentage of cells within an organoid that expressed an epitope of interest (e.g. p-S6), 2-3 images containing a minimum of 3 organoids were analyzed and the average value per animal was reported. To establish the number of hair cells within cochlear explants confocal z-stacks spanning the supporting cell and hair cell layer were exported into Photoshop CS6 (Adobe). Existing (MYO7A^+^) and newly formed hair cells (MYO7A^+^ SOX2^+^) were manually counted. The length of the analyzed cochlear segment was measure using ImageJ (https://imagej.nih.gov/ij/)). At a minimum 2 independent experiments were conducted in which at a minimum 3 cochlear explants/ organoid cultures per genotype and treatment were analyzed and reported.

### RNA extraction and RT-qPCR

Total RNA was extracted using the miRNeasy Micro Kit (Qiagen, no. 217084). The mRNA was reverse transcribed into cDNA using the iScript cDNA synthesis kit (Bio-Rad, no. 1708889). SYBR Green-based qPCR was performed by using Fast SYBR Green Master Mix reagent (Thermo Fisher Scientific, no. 4385612) and gene-specific primers. QPCRs were carried out in triplicate using a CFX Real-Time PCR System (Bio-Rad). *Rpl19* was used as an endogenous reference gene. Relative gene expression was calculated using the ΔΔCT method (91). QPCR primers are listed in Table S2.

### Immunoblotting

Cochlear epithelia or organoids were lysed with RIPA buffer (Sigma-Aldrich, no. R0278) supplemented with protease inhibitor (Sigma-Aldrich, no.11697498001), phosphatase Inhibitor cocktail 2 (Sigma-Aldrich, no. P5726) and phosphatase inhibitor cocktail 3 (Sigma-Aldrich, no. P0044). Equal amounts of protein lysate were resolved on NuPAGE 4-12% Bis-Tris Gels (ThermoFisher, no. NP0321PK2) and transferred to a PVDF membrane (Bio-Rad) by electrophoresis. Membranes were blocked with 5% (w/vol) nonfat dry milk in TBST and immunoblotted with primary antibodies and HRP-conjugated secondary antibodies (Jackson Immuno Research) according to manufacturer’s instructions. This was followed by washes in TBST and incubation in SuperSignal™ West Pico PLUS Chemiluminescent Substrate (ThermoFisher, no.34580). The resulting chemiluminescence was captured using X-ray films. The following primary antibodies were used: rabbit anti-Akt (Cell Signaling, no. 4691, 1:1000), rabbit anti-p-Akt (Ser473) (Cell Signaling, no. 4060, 1:1000), rabbit anti-p-4E-BP1 (Thr37/46) (Cell Signaling, no. 2855, 1:1000), rabbit anti-4E-BP1 (Cell Signaling, no. 9644, 1:2000), rabbit anti-human LIN28B (Cell Signaling, no. 4196, 1:2000), rabbit anti-mouse LIN28B (Cell Signaling, no. 5422, 1:500), rabbit anti-p-S6 (Ser240/244) (Cell Signaling, no. 5364, 1:500), mouse anti-β-actin (Santa Cruz Biotechnology, no. sc-47778, 1:500). To quantify protein levels, X-ray films were scanned, digital images converted to grey scale and the relative density (intensity) of bands was analyzed using the Gel Analysis plug in ImageJ (https://imagej.nih.gov/ij/).

### Statistical Analysis

All results were confirmed by at least 2 independent experiments. The sample size (n) represents the number of animals analyzed per group. Animals (biological replicates) were allocated into control or experimental groups based on genotype and/or type of treatment. To avoid bias, masking was used during data analysis. Data was analyzed using Graphpad Prism 8.0 (Graphpad Software Inc., La Jolla, CA). Relevant information for each experiment including sample size, statistical tests and reported p-values are found in the legend corresponding to each figure.

## ACKNOWLEDGMENTS

We thank the members of the Doetzlhofer Laboratory for the help and advice provided throughout the course of this study. We thank Dr. Georg Dailey for essential mouse reagents. The work was supported by NIDCD Grants DC011571 (A.D.), DC005211 (Sensory Mechanisms Research Core Center) and David M. Rubenstein Fund for Hearing Research (A.D.).

## CITATIONS

1. M. E. Warchol, Sensory regeneration in the vertebrate inner ear: differences at the levels of cells and species. Hear Res 273, 72–79 (2011).

2. D. M. Fekete, S. Muthukumar, D. Karagogeos, Hair cells and supporting cells share a common progenitor in the avian inner ear. J Neurosci 18, 7811–7821 (1998).

3. E. C. Driver, L. Sillers, T. M. Coate, M. F. Rose, M. W. Kelley, The Atoh1-lineage gives rise to hair cells and supporting cells within the mammalian cochlea. Dev Biol 376, 86–98 (2013).

4. A. E. Kiernan et al., Sox2 is required for sensory organ development in the mammalian inner ear. Nature 434, 1031–1035 (2005).

5. M. Ahmed et al., Eya1-Six1 interaction is sufficient to induce hair cell fate in the cochlea by activating Atoh1 expression in cooperation with Sox2. Dev Cell 22, 377–390 (2012).

6. J. S. Kempfle, J. L. Turban, A. S. Edge, Sox2 in the differentiation of cochlear progenitor cells. Sci Rep 6, 23293 (2016).

7. N. A. Bermingham et al., Math1: an essential gene for the generation of inner ear hair cells. Science 284, 1837–1841 (1999).

8. R. M. Lewis, C. R. Hume, J. S. Stone, Atoh1 expression and function during auditory hair cell regeneration in post-hatch chickens. Hear Res 289, 74–85 (2012).

9. K. L. Hicks, S. R. Wisner, B. C. Cox, J. S. Stone, Atoh1 is required in supporting cells for regeneration of vestibular hair cells in adult mice. Hear Res 385, 107838 (2020).

10. D. W. Roberson, J. A. Alosi, D. A. Cotanche, Direct transdifferentiation gives rise to the earliest new hair cells in regenerating avian auditory epithelium. J Neurosci Res 78, 461–471 (2004).

11. J. Shang, J. Cafaro, R. Nehmer, J. Stone, Supporting cell division is not required for regeneration of auditory hair cells after ototoxic injury in vitro. J Assoc Res Otolaryngol 11, 203–222 (2010).

12. J. T. Corwin, D. A. Cotanche, Regeneration of sensory hair cells after acoustic trauma. Science 240, 1772–1774 (1988).

13. B. M. Ryals, E. W. Rubel, Hair cell regeneration after acoustic trauma in adult Coturnix quail. Science 240, 1774–1776 (1988).

14. B. C. Cox et al., Spontaneous hair cell regeneration in the neonatal mouse cochlea in vivo. Development 141, 816–829 (2014).

15. L. Hu et al., Diphtheria Toxin-Induced Cell Death Triggers Wnt-Dependent Hair Cell Regeneration in Neonatal Mice. J Neurosci 36, 9479–9489 (2016).

16. S. Korrapati, I. Roux, E. Glowatzki, A. Doetzlhofer, Notch signaling limits supporting cell plasticity in the hair cell-damaged early postnatal murine cochlea. PLoS One 8, e73276 (2013).

17. K. Mizutari et al., Notch inhibition induces cochlear hair cell regeneration and recovery of hearing after acoustic trauma. Neuron 77, 58–69 (2013).

18. W. Li et al., Notch inhibition induces mitotically generated hair cells in mammalian cochleae via activating the Wnt pathway. Proc Natl Acad Sci U S A 112, 166–171 (2015).

19. R. Chai et al., Wnt signaling induces proliferation of sensory precursors in the postnatal mouse cochlea. Proc Natl Acad Sci U S A 109, 8167–8172 (2012).

20. F. Shi, L. Hu, A. S. Edge, Generation of hair cells in neonatal mice by beta-catenin overexpression in Lgr5-positive cochlear progenitors. Proc Natl Acad Sci U S A 110, 13851–13856 (2013).

21. B. R. Kuo, E. M. Baldwin, W. S. Layman, M. M. Taketo, J. Zuo, In Vivo Cochlear Hair Cell Generation and Survival by Coactivation of beta-Catenin and Atoh1. J Neurosci 35, 10786–10798 (2015).

22. W. Ni et al., Wnt activation followed by Notch inhibition promotes mitotic hair cell regeneration in the postnatal mouse cochlea. Oncotarget 7, 66754–66768 (2016).

23. K. B. Szarama, N. Gavara, R. S. Petralia, M. W. Kelley, R. S. Chadwick, Cytoskeletal changes in actin and microtubules underlie the developing surface mechanical properties of sensory and supporting cells in the mouse cochlea. Development 139, 2187–2197 (2012).

24. K. Legendre, S. Safieddine, P. Kussel-Andermann, C. Petit, A. El-Amraoui, alphaII-betaV spectrin bridges the plasma membrane and cortical lattice in the lateral wall of the auditory outer hair cells. J Cell Sci 121, 3347–3356 (2008).

25. A. Lelli, Y. Asai, A. Forge, J. R. Holt, G. S. Geleoc, Tonotopic gradient in the developmental acquisition of sensory transduction in outer hair cells of the mouse cochlea. J Neurophysiol 101, 2961–2973 (2009).

26. J. C. Maass et al., Changes in the regulation of the Notch signaling pathway are temporally correlated with regenerative failure in the mouse cochlea. Front Cell Neurosci 9, 110 (2015).

27. A. Samarajeewa et al., Transcriptional response to Wnt activation regulates the regenerative capacity of the mammalian cochlea. Development 145 (2018).

28. E. J. Golden, A. Benito-Gonzalez, A. Doetzlhofer, The RNA-binding protein LIN28B regulates developmental timing in the mammalian cochlea. Proc Natl Acad Sci U S A 112, E3864–3873 (2015).

29. V. Ambros, H. R. Horvitz, Heterochronic mutants of the nematode Caenorhabditis elegans. Science 226, 409–416 (1984).

30. B. J. Reinhart et al., The 21-nucleotide let-7 RNA regulates developmental timing in Caenorhabditis elegans. Nature 403, 901–906 (2000).

31. F. Rehfeld, A. M. Rohde, D. T. Nguyen, F. G. Wulczyn, Lin28 and let-7: ancient milestones on the road from pluripotency to neurogenesis. Cell Tissue Res 359, 145–160 (2015).

32. N. Shyh-Chang, G. Q. Daley, Lin28: primal regulator of growth and metabolism in stem cells. Cell Stem Cell 12, 395–406 (2013).

33. P. M. White, A. Doetzlhofer, Y. S. Lee, A. K. Groves, N. Segil, Mammalian cochlear supporting cells can divide and trans-differentiate into hair cells. Nature 441, 984–987 (2006).

34. K. Oshima et al., Differential distribution of stem cells in the auditory and vestibular organs of the inner ear. J Assoc Res Otolaryngol 8, 18–31 (2007).

35. W. J. McLean et al., Clonal Expansion of Lgr5-Positive Cells from Mammalian Cochlea and High-Purity Generation of Sensory Hair Cells. Cell Rep 18, 1917–1929 (2017).

36. J. Waldhaus, R. Durruthy-Durruthy, S. Heller, Quantitative High-Resolution Cellular Map of the Organ of Corti. Cell Rep 11, 1385–1399 (2015).

37. E. A. Lumpkin et al., Math1-driven GFP expression in the developing nervous system of transgenic mice. Gene Expr Patterns 3, 389–395 (2003).

38. Z. P. Stojanova, T. Kwan, N. Segil, Epigenetic regulation of Atoh1 guides hair cell development in the mammalian cochlea. Development 143, 1632 (2016).

39. H. Zhu et al., The Lin28/let-7 axis regulates glucose metabolism. Cell 147, 81–94 (2011).

40. A. G. Coppens, A. Resibois, L. Poncelet, Immunolocalization of calbindin D28k and calretinin in the dog cochlea during postnatal development. Hear Res 145, 101–110 (2000).

41. N. Sakaguchi, M. T. Henzl, I. Thalmann, R. Thalmann, B. A. Schulte, Oncomodulin is expressed exclusively by outer hair cells in the organ of Corti. J Histochem Cytochem 46, 29–40 (1998).

42. K. Shim, G. Minowada, D. E. Coling, G. R. Martin, Sprouty2, a mouse deafness gene, regulates cell fate decisions in the auditory sensory epithelium by antagonizing FGF signaling. Dev Cell 8, 553–564 (2005).

43. P. J. Lanford, R. Shailam, C. R. Norton, T. Gridley, M. W. Kelley, Expression of Math1 and HES5 in the cochleae of wildtype and Jag2 mutant mice. J Assoc Res Otolaryngol 1, 161–171 (2000).

44. A. Doetzlhofer, P. White, Y. S. Lee, A. Groves, N. Segil, Prospective identification and purification of hair cell and supporting cell progenitors from the embryonic cochlea. Brain Res 1091, 282–288 (2006).

45. P. Chen, N. Segil, p27(Kip1) links cell proliferation to morphogenesis in the developing organ of Corti. Development 126, 1581–1590 (1999).

46. H. Lowenheim et al., Gene disruption of p27(Kip1) allows cell proliferation in the postnatal and adult organ of corti. Proc Natl Acad Sci U S A 96, 4084–4088 (1999).

47. Z. Liu et al., Regulation of p27Kip1 by Sox2 maintains quiescence of inner pillar cells in the murine auditory sensory epithelium. J Neurosci 32, 10530–10540 (2012).

48. J. C. Maass et al., Transcriptomic Analysis of Mouse Cochlear Supporting Cell Maturation Reveals Large-Scale Changes in Notch Responsiveness Prior to the Onset of Hearing. PLoS One 11, e0167286 (2016).

49. H. C. Wang et al., Spontaneous Activity of Cochlear Hair Cells Triggered by Fluid Secretion Mechanism in Adjacent Support Cells. Cell 163, 1348–1359 (2015).

50. E. J. Son et al., Conserved role of Sonic Hedgehog in tonotopic organization of the avian basilar papilla and mammalian cochlea. Proc Natl Acad Sci U S A 112, 3746–3751 (2015).

51. M. Holley et al., Emx2 and early hair cell development in the mouse inner ear. Dev Biol 340, 547–556 (2010).

52. H. Laine, M. Sulg, A. Kirjavainen, U. Pirvola, Cell cycle regulation in the inner ear sensory epithelia: role of cyclin D1 and cyclin-dependent kinase inhibitors. Dev Biol 337, 134–146 (2010).

53. G. Shinoda et al., Fetal deficiency of lin28 programs life-long aberrations in growth and glucose metabolism. Stem Cells 31, 1563–1573 (2013).

54. Y. Ruzankina et al., Deletion of the developmentally essential gene ATR in adult mice leads to age-related phenotypes and stem cell loss. Cell Stem Cell 1, 113–126 (2007).

55. R. A. Saxton, D. M. Sabatini, mTOR Signaling in Growth, Metabolism, and Disease. Cell 169, 361–371 (2017).

56. V. Albert, M. N. Hall, mTOR signaling in cellular and organismal energetics. Curr Opin Cell Biol 33, 55–66 (2015).

57. J. A. Hutchinson, N. P. Shanware, H. Chang, R. S. Tibbetts, Regulation of ribosomal protein S6 phosphorylation by casein kinase 1 and protein phosphatase 1. J Biol Chem 286, 8688–8696 (2011).

58. S. R. von Manteuffel et al., The insulin-induced signalling pathway leading to S6 and initiation factor 4E binding protein 1 phosphorylation bifurcates at a rapamycin-sensitive point immediately upstream of p70s6k. Mol Cell Biol 17, 5426–5436 (1997).

59. G. J. Brunn et al., Phosphorylation of the translational repressor PHAS-I by the mammalian target of rapamycin. Science 277, 99–101 (1997).

60. V. Facchinetti et al., The mammalian target of rapamycin complex 2 controls folding and stability of Akt and protein kinase C. EMBO J 27, 1932–1943 (2008).

61. D. A. Guertin, D. M. Sabatini, The pharmacology of mTOR inhibition. Sci Signal 2, pe24 (2009).

62. K. Leitmeyer et al., Inhibition of mTOR by Rapamycin Results in Auditory Hair Cell Damage and Decreased Spiral Ganglion Neuron Outgrowth and Neurite Formation In Vitro. Biomed Res Int 2015, 925890 (2015).

63. W. Ni et al., Extensive Supporting Cell Proliferation and Mitotic Hair Cell Generation by In Vivo Genetic Reprogramming in the Neonatal Mouse Cochlea. J Neurosci 36, 8734–8745 (2016).

64. J. Godwin, The promise of perfect adult tissue repair and regeneration in mammals: Learning from regenerative amphibians and fish. Bioessays 36, 861–871 (2014).

65. P. J. Atkinson et al., Sox2 haploinsufficiency primes regeneration and Wnt responsiveness in the mouse cochlea. J Clin Invest 128, 1641–1656 (2018).

66. X. Duan et al., Subtype-specific regeneration of retinal ganglion cells following axotomy: effects of osteopontin and mTOR signaling. Neuron 85, 1244–1256 (2015).

67. K. K. Park et al., Promoting axon regeneration in the adult CNS by modulation of the PTEN/mTOR pathway. Science 322, 963–966 (2008).

68. S. G. Willet et al., Regenerative proliferation of differentiated cells by mTORC1-dependent paligenosis. EMBO J 37 (2018).

69. J. T. Rodgers et al., mTORC1 controls the adaptive transition of quiescent stem cells from G0 to G(Alert). Nature 510, 393–396 (2014).

70. M. Laplante, D. M. Sabatini, mTOR signaling in growth control and disease. Cell 149, 274–293 (2012).

71. C. Gao et al., Autophagy negatively regulates Wnt signalling by promoting Dishevelled degradation. Nat Cell Biol 12, 781–790 (2010).

72. L. Vadlakonda, M. Pasupuleti, R. Pallu, Role of PI3K-AKT-mTOR and Wnt Signaling Pathways in Transition of G1-S Phase of Cell Cycle in Cancer Cells. Front Oncol 3, 85 (2013).

73. M. Yang et al., Lin28 promotes the proliferative capacity of neural progenitor cells in brain development. Development 142, 1616–1627 (2015).

74. A. N. Dubinsky et al., Let-7 coordinately suppresses components of the amino acid sensing pathway to repress mTORC1 and induce autophagy. Cell Metab 20, 626–638 (2014).

75. A. Polesskaya et al., Lin-28 binds IGF-2 mRNA and participates in skeletal myogenesis by increasing translation efficiency. Genes Dev 21, 1125–1138 (2007).

76. S. Peng et al., Genome-wide studies reveal that Lin28 enhances the translation of genes important for growth and survival of human embryonic stem cells. Stem Cells 29, 496–504 (2011).

77. J. Nishino, S. Kim, Y. Zhu, H. Zhu, S. J. Morrison, A network of heterochronic genes including Imp1 regulates temporal changes in stem cell properties. Elife 2, e00924 (2013).

78. Y. C. Lin et al., Human TRIM71 and its nematode homologue are targets of let-7 microRNA and its zebrafish orthologue is essential for development. Mol Biol Evol 24, 2525–2534 (2007).

79. Y. S. Lee, A. Dutta, The tumor suppressor microRNA let-7 represses the HMGA2 oncogene. Genes Dev 21, 1025–1030 (2007).

80. N. Shyh-Chang et al., Lin28 enhances tissue repair by reprogramming cellular metabolism. Cell 155, 778–792 (2013).

81. Y. Zhang et al., Lin28 enhances de novo fatty acid synthesis to promote cancer progression via SREBP-1. EMBO Rep 20, e48115 (2019).

82. Z. N. Sayyid, T. Wang, L. Chen, S. M. Jones, A. G. Cheng, Atoh1 Directs Regeneration and Functional Recovery of the Mature Mouse Vestibular System. Cell Rep 28, 312–324.e314 (2019).

83. M. Roccio, P. Senn, S. Heller, Novel insights into inner ear development and regeneration for targeted hearing loss therapies. Hear Res 10.1016/j.heares.2019.107859, 107859 (2019).

84. G. Wan, G. Corfas, J. S. Stone, Inner ear supporting cells: rethinking the silent majority. Semin Cell Dev Biol 24, 448–459 (2013).

85. R. Ramachandran, B. V. Fausett, D. Goldman, Ascl1a regulates Muller glia dedifferentiation and retinal regeneration through a Lin-28-dependent, let-7 microRNA signalling pathway. Nat Cell Biol 12, 1101–1107 (2010).

86. S. Kaur et al., let-7 MicroRNA-Mediated Regulation of Shh Signaling and the Gene Regulatory Network Is Essential for Retina Regeneration. Cell Rep 23, 1409–1423 (2018).

87. F. Elsaeidi et al., Notch Suppression Collaborates with Ascl1 and Lin28 to Unleash a Regenerative Response in Fish Retina, But Not in Mice. J Neurosci 38, 2246–2261 (2018).

88. K. Yao et al., Wnt Regulates Proliferation and Neurogenic Potential of Muller Glial Cells via a Lin28/let-7 miRNA-Dependent Pathway in Adult Mammalian Retinas. Cell Rep 17, 165–178 (2016).

89. S. A. Bucks et al., Supporting cells remove and replace sensory receptor hair cells in a balance organ of adult mice. Elife 6 (2017).

90. A. Doetzlhofer et al., Hey2 regulation by FGF provides a Notch-independent mechanism for maintaining pillar cell fate in the organ of Corti. Dev Cell 16, 58–69 (2009).

91. T. D. Schmittgen, K. J. Livak, Analyzing real-time PCR data by the comparative C(T) method. Nat Protoc 3, 1101–1108 (2008).

