## Supplemental Material for "LIN28B controls the regenerative capacity of neonatal murine auditory supporting cells through activation of mTOR signaling"

### SUPPLEMENTAL FIGURES

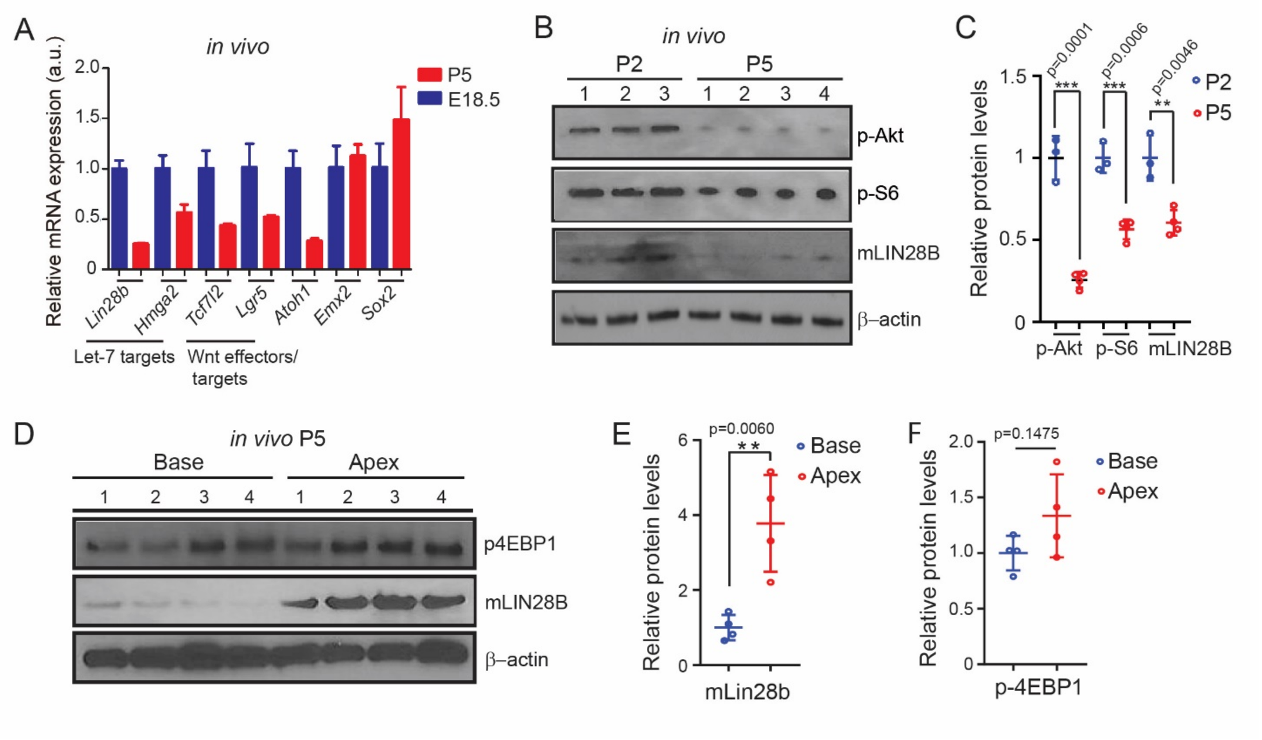


**Fig. S1.** LIN28B expression positively correlates with mTOR activity in the maturing cochlea. (A) RT-qPCR was used to analyze mRNA abundance of *let-7* targets (*Lin28b*, *Hmga2*), Wnt signaling effectors/targets (*Tcf7l2*, *Lgr5*) and *Atoh1*, *Emx2*, *Sox2* in cochlear sensory epithelia obtained from wild type mice stages E18.5 (blue) and P5 (red) (graphed are mean ± SD, technical replicate, shown representative experiment, 3 independent experiments). (B) Immunoblots for p-Akt, p-S6, murine (m) LIN28B and β-actin (loading control) using protein lysates of acutely isolated cochlear sensory epithelia from wild type mice stages P2 (n=3) and P5 (n=4). (C) Normalized p-Akt, p-S6 and murine (m) LIN28B protein expression in (B) (n=3 animals for P2 and n=4 mice for P5, from 1 representative experiment, 2 independent experiments). (D) Immunoblots for p-4EBP1, murine (m) LIN28B and β-actin (loading control) using protein lysates of acutely isolated sensory epithelia obtained from the cochlear apex and base of wild type mice stage P5. (E-F) Normalized LIN28B protein levels in (D) (n=4 animals). (F) Normalized p-4EBP1 protein levels in (D) (n=4 animals, from 1 representative experiment, 2 independent experiments). Graphed are individual data points and mean ± SD, 2-tailed, unpaired Student's t-test was used to calculate p-values in (C), (E) and (F). Abbreviation: a.u., arbitrary unit.

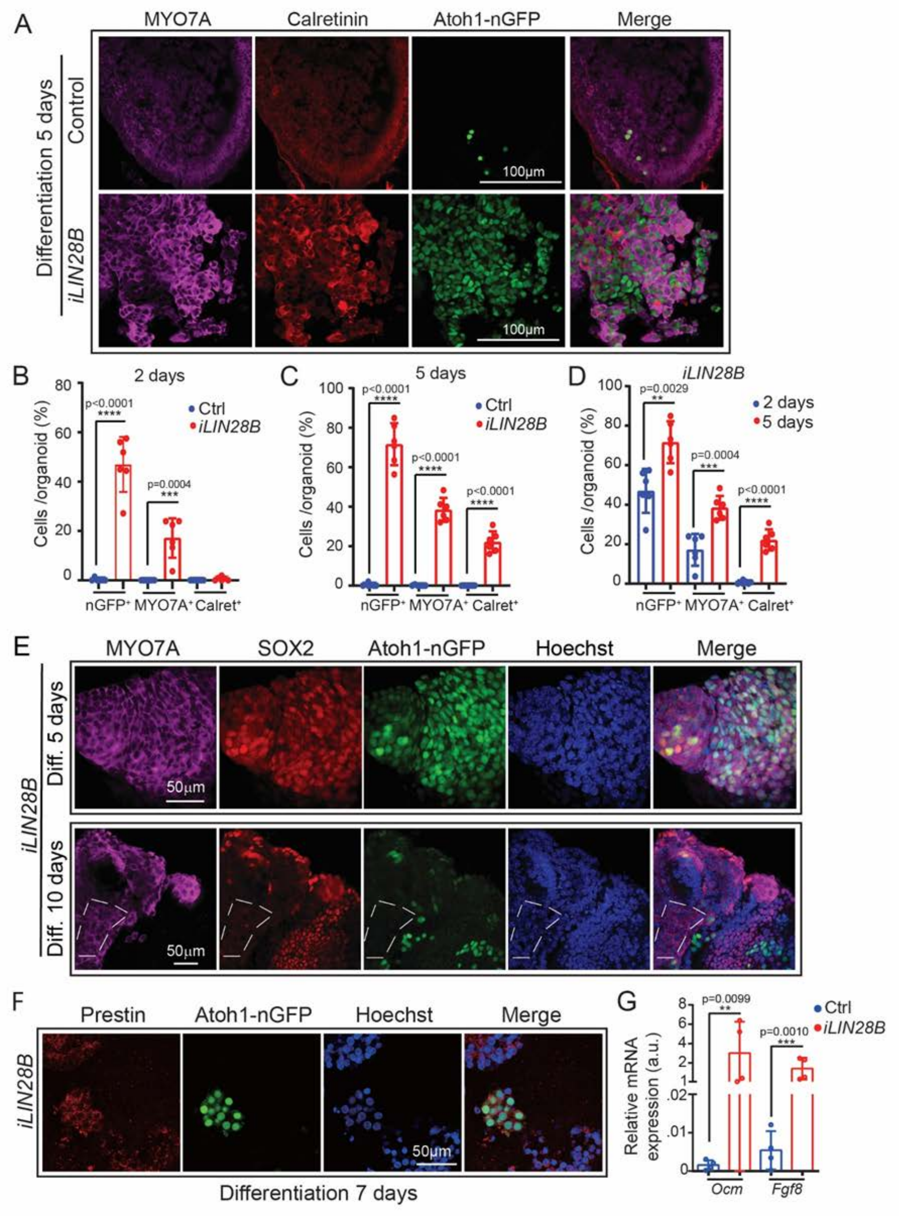


**Fig. S2.** Newly formed hair cells in LIN28B overexpressing organoids express inner and outer hair cell markers. Cochlear organoid cultures were established from stage P5 Atoh1-nGFP iLIN28B transgenic mice and Atoh1-nGFP control littermates that lacked the LIN28B transgene and were cultured as described in Fig. 2 A. 2-tailed, unpaired Student's t-test was used to calculate p-values. Note that the individual date points in (B), (C) (D) and (G) represent the average values per animal. (A) Confocal images of control and LIN28B overexpressing organoids after 2 and 5 days of differentiation. Atoh1-GFP (green) and MYO7A (magenta) marks newly formed hair cells. Calretinin (red) marks presumptive inner hair cells. (B-D) Percentage of Atoh1-nGFP, MYO7A and calretinin positive cells per organoid in control (Ctrl) and LIN28B overexpressing (iLIN28B) organoid cultures after 2 days (B and D), 5 days (C and D) of differentiation (graphed are average values for each animal and their mean ± SD, n=6 animals per group, from 2 independent experiments). (E) Confocal images of LIN28B overexpressing organoids after 5 and 10 days of differentiation. Immature hair cells are identified by their co-expression of Atoh1-GFP (green) and MYO7A (magenta) and SOX2 (red). White dashed lines encircle a group of more mature hair cells that express MYO7A but lack SOX2 and Atoh1-GFP expression. Hoechst (blue) labels cell nuclei. (F) Confocal images of control and LIN28B overexpressing organoids after 7 days of differentiation. Atoh1-nGFP (green) and prestin (red) co-expression marks presumptive outer hair cells, Hoechst (blue) labels cell nuclei. (G) RT-PCR of inner (*Fgf8*) and outer (oncomodulin, *Ocm*) hair cell-specific gene expression in control and iLIN28B transgenic organoids after 7 days of differentiation (mean ± SD, n=4 animals per group, from 2 independent experiments).

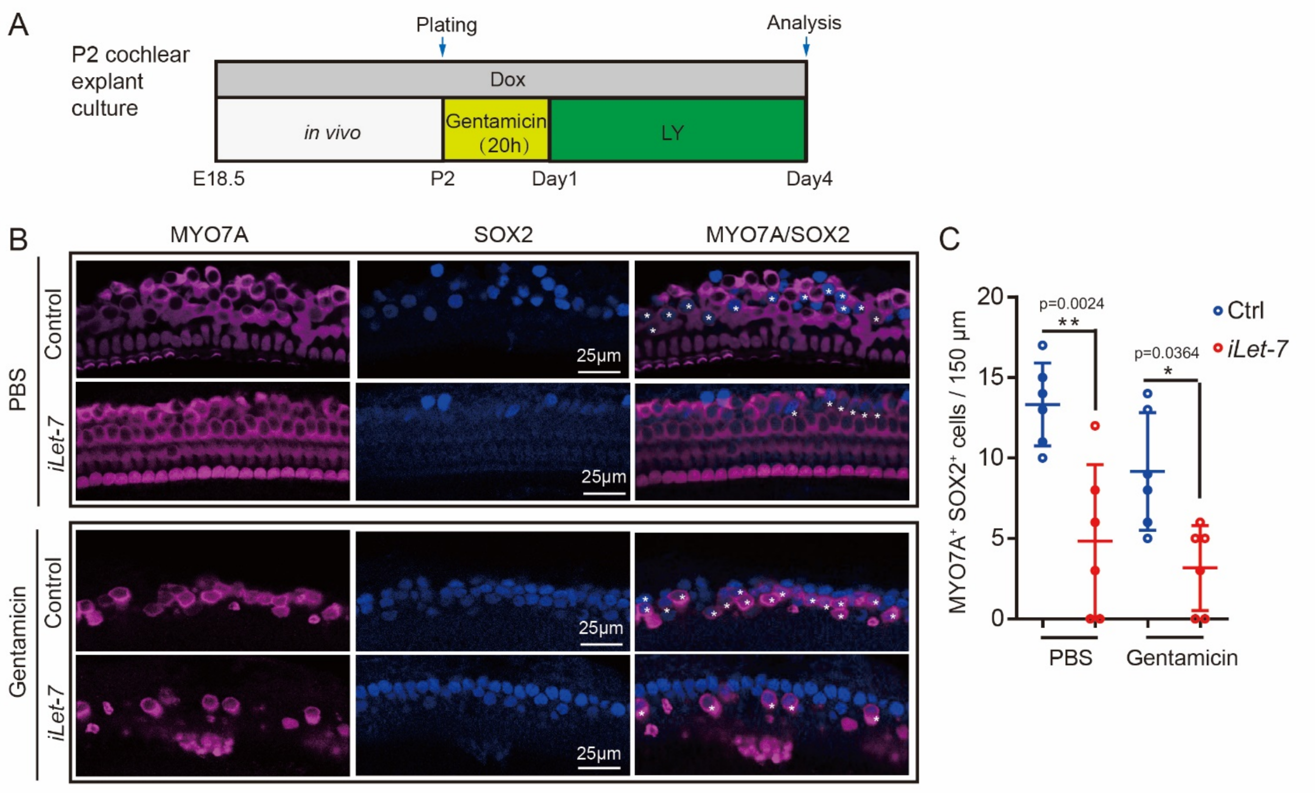


**Fig. S3.** *Let-7* overexpression inhibits hair cell regeneration in early postnatal cochlear explants. *(A)* Experimental strategy. Cochlear explant cultures were established from P2 *iLet-7* transgenic pups and littermates that lacked *let-7g* transgene (control). To ablate hair cells, one cochlea from each animal received gentamicin (100 µg/ml), while the other cochlea received PBS (vehicle control) for 20 hours. To induce supporting cell-to-hair cell conversion, all cultures were treated with Notch inhibitor LY411575 for 3 days starting at day 1. *(B)* Shown are representative confocal images of mid-apical turn of control and *let-7g* overexpressing (*iLet-7*) cochlear explants immuno-stained for MYO7A (magenta) and SOX2 (blue). Note that new hair cells express MYO7A and SOX2, whereas pre-existing hair cells only express MYO7A. *(C)* Quantification of newly formed hair cells (MYO7A+, SOX2+) in control (blue) and *let-7g* overexpressing cochlear explants (red) in *(B)*. Graphed are individual data points, representing average values per animal, and mean ± SD, n=6 mice per group, from 2 independent experiments. 2-way ANOVA with Tukey's correction was used to calculate p-values.

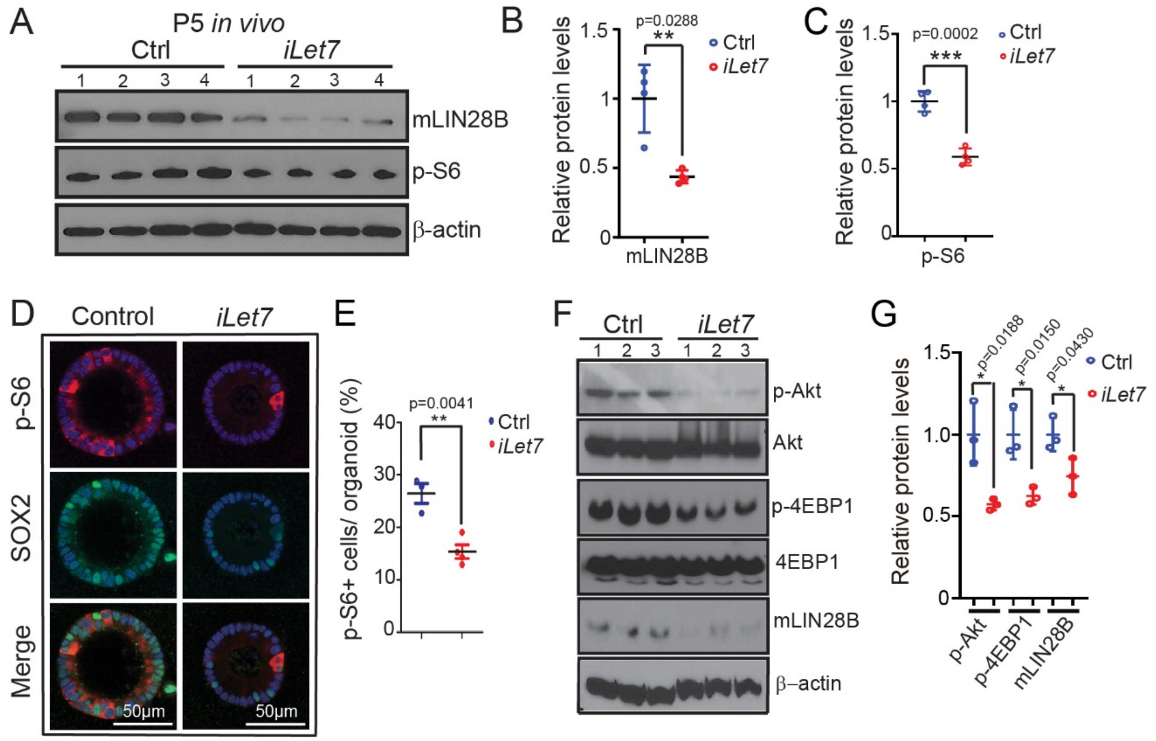


**Fig. S4.** *Let-7g* negatively regulates mTOR signaling in early postnatal cochlear epithelial cells. Graphed are individual data points and mean± SD. 2-tailed, unpaired Student's t-test was used to calculate p-values. Note that the individual data points in (*E*) represent the average values per animal. (A-C) *Let-7g* overexpression attenuates mTOR signaling in cochlear epithelial cells *in vivo*. *iLet*-7 transgenic mice and control littermates that lacked *let-7g* transgene received dox starting at E18.5 until tissue harvest at P5. *(A)* Immunoblots for p-S6, endogenous murine (m) LIN28B and β-actin using protein lysates of acutely isolated control (ctrl) and *let-7g* (*iLet-7*) overexpressing cochlear sensory epithelia. (*B-C*) Normalized murine (m) LIN28B protein expression and p-S6 protein in (*A*) (n=4 animals per group, from 1 representative experiment, 3 independent experiments). (*D-G*) *Let-7g* overexpression attenuates mTOR signaling in cochlear organoids. Cochlear organoid cultures were established from stage P2 *iLet-7* transgenic mice and control littermates. Dox was present throughout the 10-day long expansion phase. (*D*) Confocal images of control and *let-7g* overexpressing (*iLet-7*) organoids co-stained for p-S6 (red) and

SOX2 (green). Nuclei were counterstained with Hoechst (blue). (*E*) Percentage of p-S6+ cells per organoid shown in (*D*) (n=3 animals in control group and n=4 animals in *iLet-7* group, from 1 experiment). (*F*) Immunoblots for p-Akt, Akt, p-4EBP1, 4EBP1, murine (m) LIN28B and β-actin using protein lysates from control and *iLet-7* transgenic organoids. (*G*) Normalized p-Akt, p-4EBP1 and mLIN28B protein expression in (*F*) (n=3 animals per group, from 1 representative experiment, 3 independent experiments).

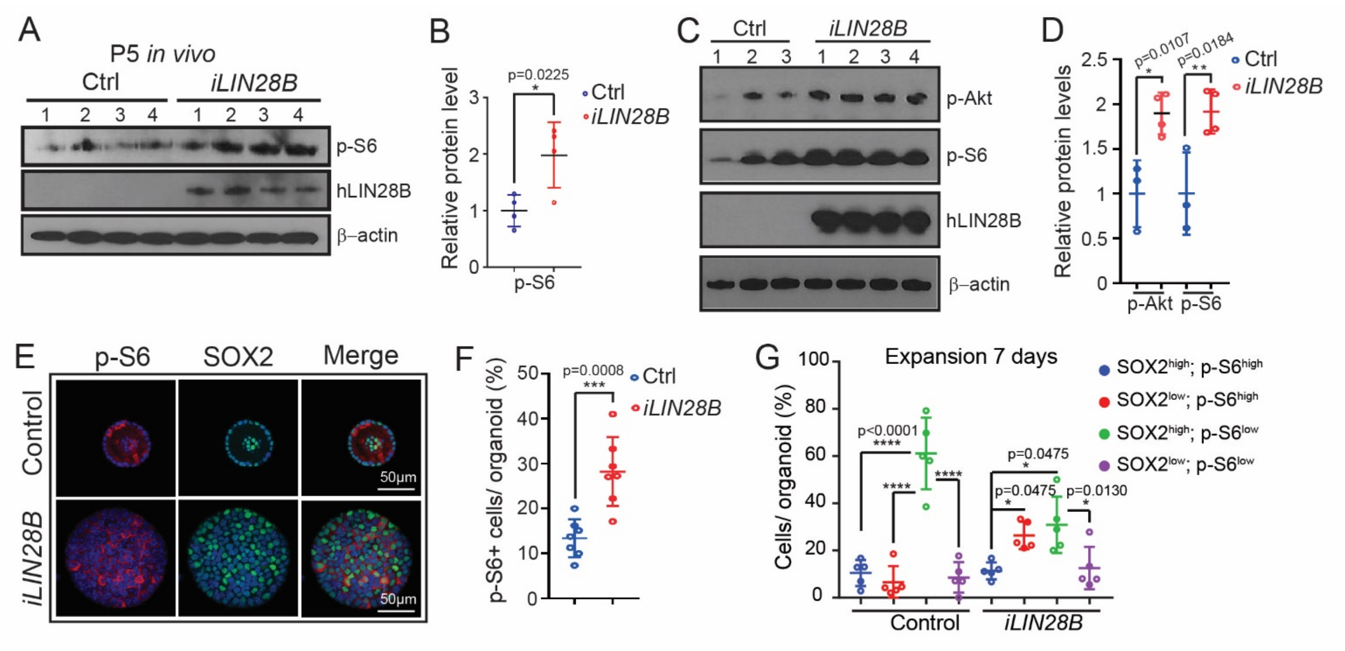


**Fig. S5.** LIN28B positively regulates mTOR signaling in early postnatal cochlear epithelial cells. (*A*-*B*) LIN28B overexpression enhances mTOR signaling in cochlear epithelial cells *in vivo*. *iLIN28B* transgenic mice and control littermates received dox starting at E18.5 until tissue harvest at P5 (A) Immunoblots for p-S6, human (h) LIN28B and β-actin using cochlear epithelial protein lysates from stage P5 control and LIN28B overexpressing mice. (B) Normalized p-S6 protein levels in (*A*) (n=4 animals per group, from 1 representative experiment, 3 independent experiments). (*C*-*G*) LIN28B overexpression increases mTOR activity in cochlear organoids. Cochlear organoid cultures were established from stage P5 iLIN28B transgenic mice and control littermates. Organoid cultures were maintained as outlined in Fig. 2A. Dox-containing culture media was replenished every other day. (*C*) Immunoblots for p-Akt, p-S6, human (h) LIN28B and β-actin using protein lysates of control and iLIN28B organoids. (*D*) Normalized p-Akt and p-S6 protein levels in (*C*) (n=3 animals in control group and n=4 animals in iLIN28B group, from 1 representative experiment, 3 independent experiments). (*E*) Confocal images of P5 control and P5 iLIN28B transgenic organoids after 9 days of expansion. Organoids were immuno-stained for mTOR target p-S6 (red) and supporting cell/pro-sensory cell marker SOX2 (green). Nuclei were counterstained with Hoechst (blue). (*F*) Percentage of p-S6+ cells per organoid shown in (*E*) (n=7 animals per group, from 2 independent experiments). (*G*) Percentage of p-S6+ cells (p-S6-high) that express SOX2 at high or low level in (*E*) (n=5 animals per group, from 2 independent experiments). Graphed are individual date points and the mean ±SD. 2-tailed, unpaired Student's t-test was used to calculate p-values. Note that the individual data points in (*F*) and (*G*) represent the average values per animal.

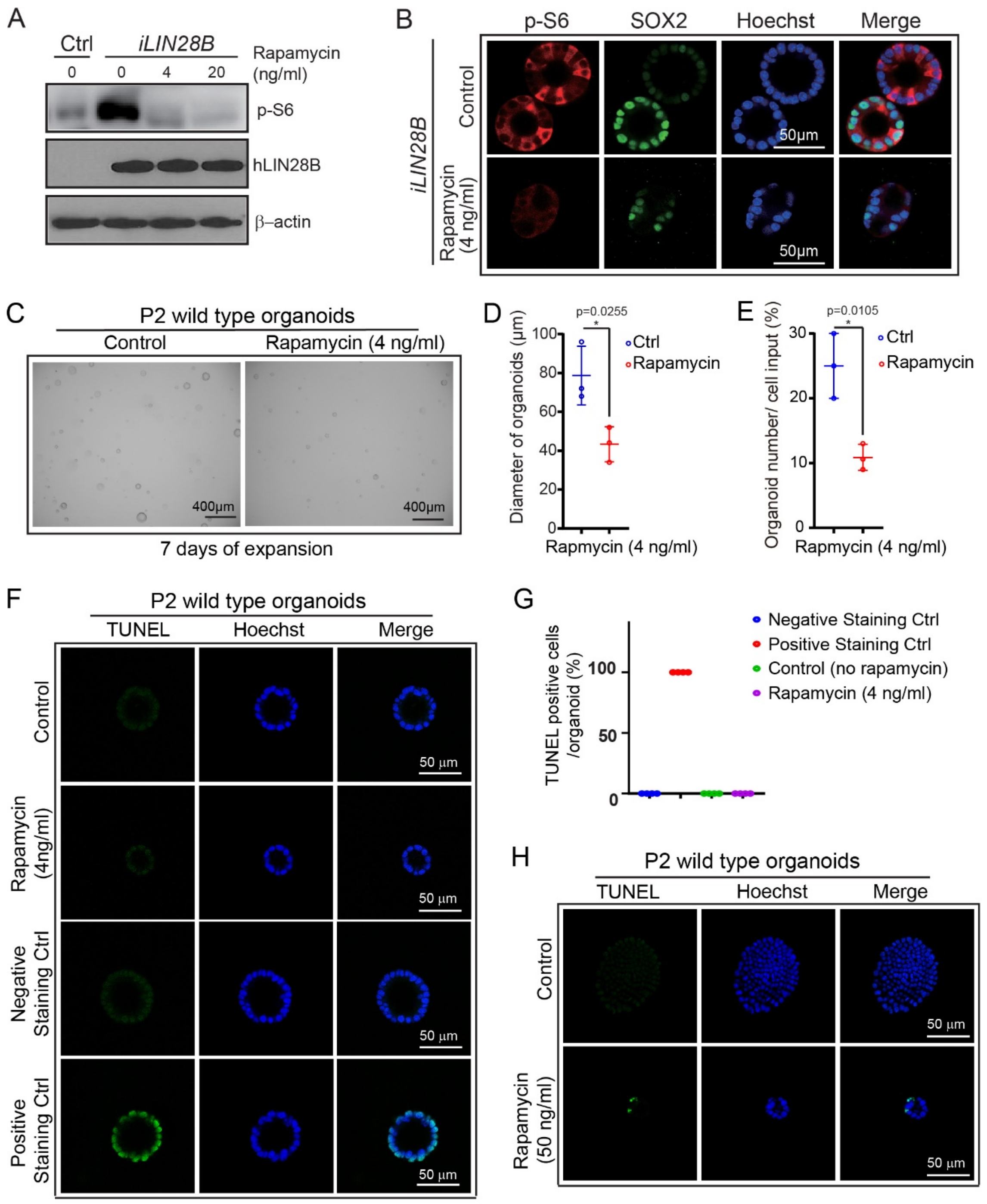


**Fig.S6.** Low dosage of rapamycin inhibits cochlear organoid growth without inducing cell death. (*A-B*) 4 ng/mL of rapamycin is effective in inhibiting mTOR activity in LIN28B overexpressing cochlear organoids. (A) Western blot of p-S6 and human (h) LIN28B in protein lysates collected from untreated (0 rapamycin) control (Ctrl) and LIN28B overexpressing (*iLIN28B*) organoids and

LIN28B overexpressing organoids that were treated for 5 hours with 4 ng/ml  or 20 ng/mL of rapamycin. Note that rapamycin treatment has no effect on LIN28B transgene expression. (*B*) Confocal images of LIN28B overexpressing organoids treated with 4 ng/mL rapamycin or vehicle control (DMSO) for 7 days. P-S6 (red) immuno-staining marks cells with high mTOR activity, SOX2 (green) marks supporting cells/ pro-sensory cells. Hoechst (blue) marks cell nuclei. (*C-H*) Cochlear organoid cultures were established from P2 wild type mice and cultured using expansion conditions (see Fig.1 A). (*C-G*) Cochlear organoids received 4 ng/mL rapamycin or vehicle control (DMSO) at day 1. Organoid growth (*C-E*) and cell death using TUNEL staining (*F-G*) was analyzed 6 days later.  Graphed are individual data points and mean ± SD, 2-tailed, unpaired Student's t-test was used to calculate p-values. (*C*) Bright field of cochlear organoids treated with 4 ng/mL rapamycin or vehicle control (DMSO). (*D*) Organoid forming efficiency and (E) organoid diameter in (*C*) (n=4 animals per group, from 2 independent experiments). (*F*) Confocal images of TUNEL (green) and Hoechst (blue) stained control (DMSO) and rapamycin (4ng/ml) treated P2 wild type organoids, including negative and positive controls for TUNEL staining. (*G*) Percentage of TUNEL+ cells in (*F*) (n=4 animals per group, from 2 independent experiments). Note that the individual data points in (*D*), (*E*) and (*G*) represent the average values per animal. (*H*) Cochlear organoids were cultured with 50 ng/mL of rapamycin or vehicle control (DMSO) starting at day 3 and cell death using TUNEL staining was analyzed 4 days later. Shown are representative confocal images of TUNEL (green) and Hoechst (blue) stained organoids.

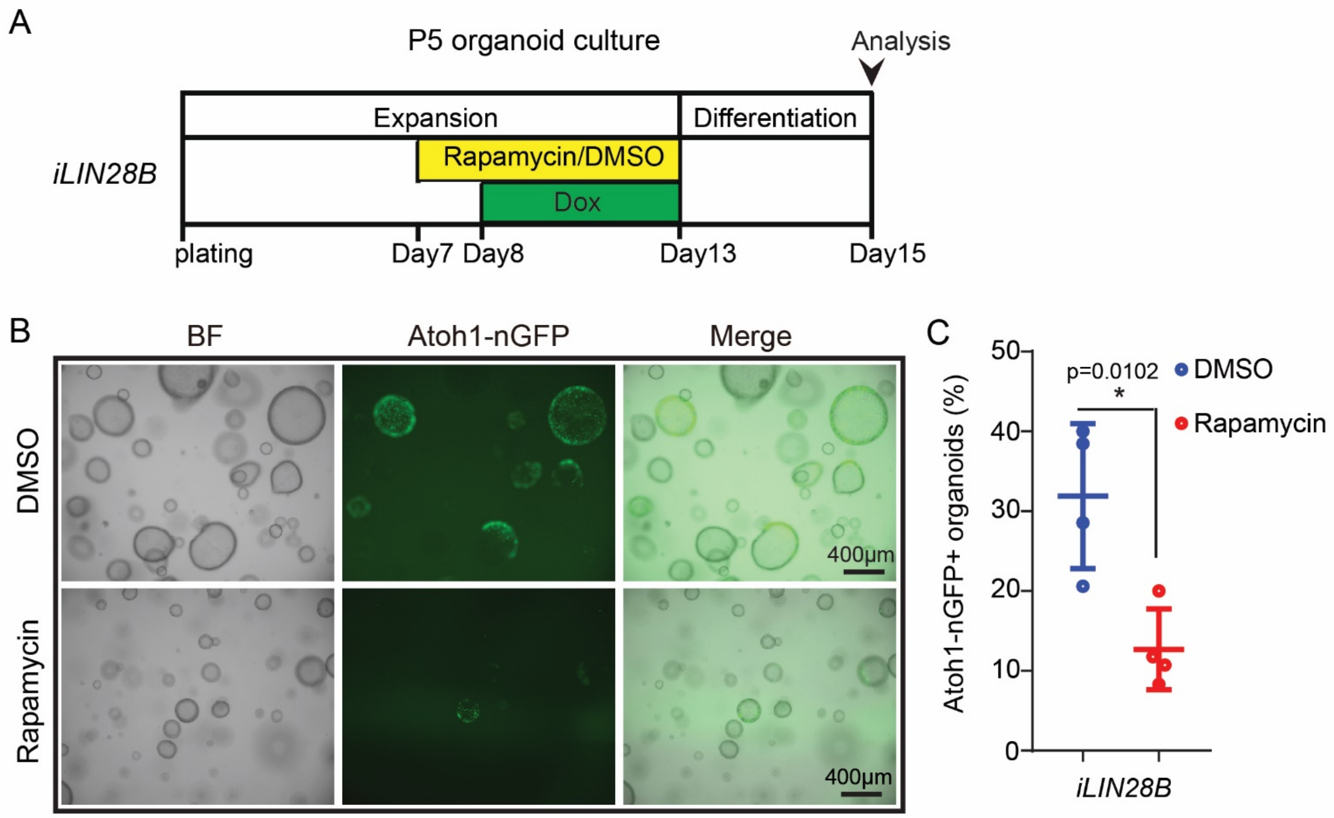


**Fig.S7.** Rapamycin pre-treatment inhibits *Atoh1* induction in LIN28B overexpressing organoids. (*A*) Experimental strategy. Organoid cultures were established from stage P5 *Atoh1-nGFP iLIN28B* transgenic mice and maintained as outlined in Fig.2 A. Rapamycin (4 ng/ml) or vehicle control (DMSO) was added to the culture media at 7 days of expansion. 1 day later, dox was added to induce LIN28B expression. The culture medium was replenished every other day. (*B*) BF and green fluorescent (Atoh1-nGFP) images of LIN28B overexpressing organoids treated with DMSO or rapamycin after 2 days of differentiation. (*C*) Percentage of Atoh1-nGFP+ organoids in (*B*) (graphed are individual data points and mean ± SD, n=4 animals per group, from 2 independent experiments). 2-tailed, unpaired Student's t-test was used to calculate p-values.

**SUPPLEMENTAL METHODS**

**Table S1.** List of genotyping primers

| **Mouse line** | **Genotyping primers** | **Product size** |
|---|---|---|
| *Atoh1-GFP, p27-GFP* | EGFP1: CGA AGG CTA CGT CCA GGA GCG CAC<br>EGFP2: GCA CGG GGC CGT CGC CGA TGG GGG TGT | TG= 300bp |
| *R26-M2rtTA* | MTR: GCG AAG AGT TTG TCC TCA ACC<br>F: AAA GTC GCT CTG AGT TGT TAT<br>WTR: GGA GCG GGA GAA ATG GAT ATG | WT=650bp<br>MT=340bp |
| *Col1A1* (*LIN28B and let-7g*) | ColA: GCA CAG CAT TGC GGA CAT GC<br>ColB: CCC TCC ATG TGT GAC CAA GG<br>ColC: GCA GAA GCG CGG CCG TCT GG | WT=300bp<br>TG=450bp |
| *UBC-CreERT2* | MT-F: GAC GTC ACC CGT TCT GTT G<br>MT-R: AGG CAA ATT TTG GTG TAC GG<br>WT-F: CTA GGC CAC AGA ATT GAA AGA TCT<br>WT-R: GTA GGT GGA AAT TCT AGC ATC ATC C | WT=324bp<br>TG=475bp |
| *Lin28a floxed* | Flox-F: TCC AAC CAG CAG TTT GCA G<br>Flox-R: GCA GCT GGT AAG AAG AAA CCT G | WT=356bp<br>Flox=500bp |
| *Lin28b floxed* | Flox-F: AAC GCA CAT TGC AAA TAC CC<br>Flox-R: TTC ATC TGG CTC CTT TCT CG | WT=221bp<br>Flox=338bp |

**Table S2.** List of qPCR primers

| Gene | Forward Primer | Reveres Primer |
|---|---|---|
| Ano1 | TTC CCT CTG GCT CCA CTC TTC | GGC ATC CAG GCG GAT CT |
| Atoh1 | ATG CAC GGG CTG AAC CA | TCG TTG TTG AAG GAC GGG ATA |
| Ccnd1 | CAA GTG TGA CCC GGA CTG C | TTG ACT CCA GAA GGG CTT CAA |
| Cybrd1 | AGA CTG CCA TGG ACC TGG AA | CCG GCA TGG ATG GAT TTC |
| Emx2 | GAA TCC GCT TTG GCT TTC TG | GAC ACA AGT CCC GAG AGT TTC C |
| F2rl1 | CGG ACC GAG AAC CTT GCA CCG | GTG AGG ATG GAC GCA GAG AACT |
| Fat3 | CAC AGC CCT TGA ATA CAG TGA | TGC CTT TGC ATC TCC TTC CT |
| Fgf8 | ATC AAC GCC ATG GCA GAA G | AGT ATC GGT CTC CAC AAT GAG CTT |
| Fst | GAA AAC CTA CCG CAA CGA ATG | TCC GGC TGC TCT TTG CAT |
| Hmga2 | CAG AAG AAA GCA GAG ACC ATT | TTG TTG TGG CCA TTT CCT AGG T |
| Lgr5 | CCC CAA TGC GTT TTC TAC GT | GAA GGA CGA CAG GAG ATT GGA T |
| Lin28a | TCC AAA GGA GAC AGG TGC TAC A | TTG CAT TCC TTG GCA TGA TG |
| Lin28b | CAT GGC ACT GGC CAC TGT AA | ATC ATG GAG ATG AAT CCG AAT CC |
| Myo7a | CCC CCT CTG AGA AGT TCG TTA A | TGT GTC CGA GTT CCG TTG AC |
| Ocm | ACC AGA GTG GAT ACC TGG ATG | CGT CGC TCT GGA ACC TCT GT |
| Pou4f3 | GCA CCA TCT GCA GGT TCG A | CCG GCT TGA GAG CGA TCA T |
| S100a1 | TGG ATG TCC AGA AGG ATG CA | CCG TTT TCA TCC AGT TCC TTC A |
| Sox2 | CCA GCG CAT GGA CAG CTA | GCT GCT CCT GCA TCA TGC T |
| Tcf7l2 | AAA CCC TCA AGG ATG CTC GTT | CCA CCG GTA CTT TGT TCG AAA |
| Trim71 | ATC GGG AGT GTG AGC TGT TG | GGC GTG AAC ATA ATG CGG TC |
| Rpl19 | GGT CTG GTT GGA TCC CAA | TGC CCG GGA ATG GAC AGT CA |
